# Multiplexed *in situ* hybridization reveals distinct lineage identities for major and minor vein initiation during maize leaf development

**DOI:** 10.1101/2024.02.05.578898

**Authors:** Chiara Perico, Maricris Zaidem, Olga Sedelnikova, Samik Bhattacharya, Christian Korfhage, Jane A. Langdale

## Abstract

Leaves of flowering plants are characterised by diverse venation patterns. Patterning begins with the selection of vein-forming procambial initial cells from within the ground meristem of a developing leaf, a process which is considered to be auxin-dependent, and continues until veins are anatomically differentiated with functional xylem and phloem. At present, the mechanisms responsible for leaf venation patterning are primarily characterized in the model eudicot *Arabidopsis thaliana* which displays a reticulate venation network. However, evidence suggests that vein development may proceed via a different mechanism in monocot leaves where venation patterning is parallel. Here, we employed Molecular Cartography, a multiplexed *in situ* hybridization technique, to analyse the spatiotemporal localisation of a subset of auxin related genes and candidate regulators of vein patterning in maize leaves. We show how different combinations of auxin influx and efflux transporters are recruited during leaf and vein specification, and how major and minor vein ranks develop with distinct identities. The localisation of the procambial marker *PIN1a* and the spatial arrangement of procambial initial cells that give rise to major and minor vein ranks further suggests that vein spacing is pre-patterned across the medio-lateral leaf axis prior to accumulation of the PIN1a auxin transporter. In contrast, patterning in the adaxial-abaxial axis occurs progressively, with markers of xylem and phloem gradually becoming polarised as differentiation proceeds. Collectively our data suggest that both lineage- and position-based mechanisms may underpin vein patterning in maize leaves.

**SIGNIFICANCE STATEMENT:** During the development of multicellular organisms specialized cell-types differentiate from pluripotent stem cells, with cell identity acquired via lineage- or position-based mechanisms. In plants, most organs develop post-embryogenesis and as such developmental processes are influenced by the external environment. To adapt to different environmental contexts and yet still form recognizable structures, position-based differentiation mechanisms are deployed in which cells adopt a certain fate depending on the activity of neighbouring cells. Such is the prevalence of position-based mechanisms in plant development that a role for lineage is rarely contemplated. Here we show that stem cells which give rise to different vein types in maize leaves are transcriptionally distinct, possibly reflecting a role for lineage-based mechanisms in the differentiation of leaf veins.

## INTRODUCTION

Within flowering plants distinct leaf venation patterns characterise the leaves of eudicotyledonous and monocotyledonous species, which display reticulate networks and parallel venation respectively (reviewed in (1)). The characteristics of parallel vein formation have been studied primarily in leaves of the monocot grass *Zea mays* (maize). As in all flowering plants, maize leaves emerge from the flanks of the shoot apical meristem (SAM) in a regular pattern. The SAM itself comprises two layers, an outer (epidermal) L1 layer and an inner L2 layer, both of which are recruited into new leaf primordia as they are formed (2). Leaves emerge sequentially on opposite sides of the SAM (alternate phyllotaxis) and at regular time intervals called plastochrons (P) (3) The incipient leaf, not yet distinguishable anatomically but for which the position is already defined is referred to as P0, the youngest visible leaf primordium as P1, and older leaves with increasing plastochron numbers (4). Primordia inception is marked by an auxin maximum and by down-regulation of the homeobox gene *KNOTTED1* (*KN1*) which acts in the SAM to promote indeterminacy within both the central zone (CZ) of stem cells and the flanking peripheral zone (PZ) that gives rise to leaves. Before primordia develop, proximo-distal (base-tip), medio-lateral (centre-margin) and adaxial-abaxial (upper-lower) leaf axes are established, each of which provides the spatial context in which veins develop (reviewed in (5)). Of particular importance is the establishment of a central layer of ground tissue in the adaxial-abaxial axis at P1, from within which all veins are subsequently initiated (6, 7). Parallel venation in maize leaves thus results from the specification of veins at regular intervals across the medio-lateral leaf axis, within a single layer of ground tissue in the adaxial-abaxial axis.

The maize leaf is characterised by several vein ranks which develop in a spatiotemporally regulated manner at successive plastochrons (Fig. 1A-C). Regardless of rank, development begins with the specification of a procambial initial cell (referred to as a pre-procambial cell in eudicots) within the central layer of the ground meristem. Once specified, this cell divides transversely to establish and extend the procambial strand in the proximo-distal leaf axis, and longitudinally/obliquely to form the procambial centre from which tissues of the vein differentiate in the medio-lateral and adaxial-abaxial leaf axes (reviewed in (8)). Major veins, which include the mid- and lateral veins, develop acropetally (base-to-tip) along the proximo-distal axis with the midvein positioned in the middle of the developing primordium by P1 and lateral veins specified either side of it between P2 and P3 (Fig. 1B-C). Minor veins, which include rank-1 and rank-2 intermediates, develop basipetally (tip-to-base) from P3 (Fig. 1B-C) and are spaced regularly between the major veins such that each vein pair is separated by four cells (two mesophyll and two bundle sheath) across the medio-lateral axis (9–11). During the final stages of vein development, differentiation occurs in the adaxial-abaxial leaf axis, positioning xylem adaxially, phloem abaxially and sclerenchyma between the vein and epidermis. The final vein rank to form develops transversely, connecting the longitudinal veins across the medio-lateral leaf axis in a striate pattern.

**Figure. 1:**
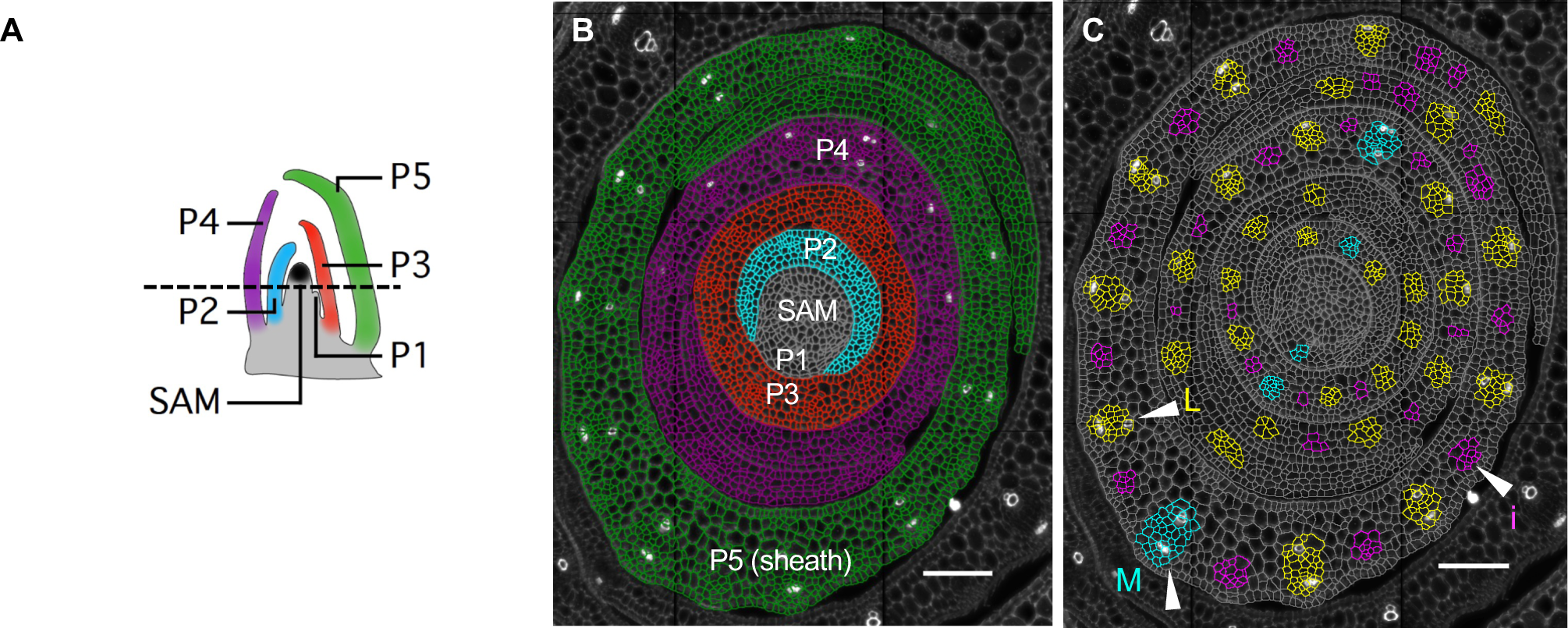
Stages of leaf and vein development in maize shoot apices. **A)** Cartoon representation of a longitudinal section across a maize shoot apex. SAM (black) indicates the Shoot Apical Meristem; younger to older stages of leaf development (leaf primordia, P) are indicated using increasing numbers and different colours. The black dotted line indicates the position of the transverse section depicted in (B). **B)** Representative transverse section of a maize shoot stained with Calcofluor and segmented with CellPose 2.0. SAM and developing leaf primordia are segmented and colour-coded as in (A). **C)** Vein ranks appear sequentially during leaf primordium development. P1: midvein only (M, cyan); P2: midvein and laterals (L, yellow); P3, P4 and P5: midvein, laterals and intermediate veins (i, magenta). Scale bars = 100 μm.

Vein patterning mechanisms have been primarily studied in the model eudicot *Arabidopsis thaliana* where the auxin efflux carrier PIN1 is required for leaf initiation on the flanks of the SAM, specification of pre-procambial cells within the leaf primordium and directional extension of the procambial strands (12). In the SAM, the simultaneous action of *PIN1* and auxin influx carriers of the *AUX1/LAX* family generates the auxin maxima that determine the position of new leaves (13–15) and in the leaf *PIN1* activity induces the auxin activated transcription factors *MONOPTEROS* (*MP/ARF5*) and *ATHB8*, which commit cells from the ground meristem to pre-procambial and then procambial fate. A feedback loop between auxin transport, MP and ATHB8 then restricts the procambial strand to a couple of cell files in width (reviewed in (16)). The reticulate vein network results from the sequential specification of vein ranks with new veins developing towards pre-existing veins, which are used as positional landmarks towards which the auxin flux is directed (12, 17). The presence of parallel veins that are precisely spaced across the medio-lateral leaf axis and a considerable expansion in the PIN and LAX transporter gene families (18) suggest that mechanisms underpinning vein patterning in monocots could be different from those found in *Arabidopsis*.

Recent genetic and transcriptome analyses have identified candidate regulators of venation patterns in monocot leaves. For example, a role for the *AINTEGUMENTA* (*ANT*) family in vein formation is supported by the finding that maize *ANT4* (19) is differentially expressed in median ground meristem (mGM) cells when compared to pre-mesophyll cells, and mutations in the *Setaria viridis AINTEGUMENTA1* (*ANT1*) gene lead to smaller leaves with distorted Kranz anatomy (20). Double mutations in *SCARECROW1* (*SCR1*) and its homeolog *SCR1h* condition narrower spaces between veins (21) and quadruple mutants with *NAKED ENDOSPERM1* (*NKD1*) and *NKD2* further enhance the spacing defect (22). In contrast, mutations in *SHORTROOT1* (*SHR1*) and its paralog *SHR2* disrupt the specification and/or subsequent development of veins (23–25), and mutations in *TOO MANY LATERALS* (*TML1*) form lateral veins in the position where the rank 1 intermediate veins that extend from blade to sheath normally develop (26). Whereas each of the genes mentioned thus far encode transcription factors, leaf venation defects have also been observed in mutants that perturb auxin biosynthesis, and transcriptome analyses have predicted a role for auxin transporters. For example, in *SPARSE INFLORESCENCE1* (*SPI1*) and *VANISHING TASSEL2* (*VT2*) mutants, reduced levels of auxin are associated with the formation of more intermediate veins in wider than normal leaves (27). Since neither pharmacological nor genetic approaches have disrupted the correlation between vein number and leaf width in monocots, any change of vein patterning accompanied by a change in leaf width is unlikely to result from directly disrupting the patterning mechanism (28). However, expression of the *AtPIN1* ortholog *ZmPIN1a* in all vein ranks from early stages of vein and leaf development (27) and a similar spatial expression pattern of the *LAX1* (note named *LAX2* in (19)) and *LAX2* (29) influx carriers at later stages of vein development, strongly suggest that auxin flux plays a role in vein patterning in the maize leaf. Although these observations are too fragmented to reveal any form of gene regulatory network, consistent expression profiles and mutant phenotypes of orthologs in different grass species (e.g. *TML1*, *SHR1, PIN1a/PIN1d*) suggest that core components of the mechanism underpinning parallel venation patterns in monocots have been identified.

Here we employed the Molecular Cartography platform to investigate cellular relationships during early stages of maize leaf development. Molecular Cartography is a high-throughput quantitative *in situ* hybridization technology that combines single cell detection of transcripts with a visual interface to identify spatial localisation within a tissue (30–33). Focusing on candidate regulators of leaf initiation and vein patterning, we qualitatively and quantitatively defined spatial and temporal patterns of transcript accumulation across consecutive stages of development in maize shoot apices. These analyses revealed developmental patterns not previously detected by either morphological or cell lineage analyses, providing insight into the specification of procambial initial cells, the establishment of procambial centres giving rise to different vein ranks, and the differentiation of tissues within each vein.

## RESULTS

### Molecular Cartography reliably reports transcript accumulation patterns in maize shoot apices

We sought to generate a spatial expression atlas in maize shoot apices for genes in which an aberrant leaf venation phenotype has been described previously in either monocots or dicots (Table S1). This group of genes included auxin efflux and influx transporters such as *PIN* and LAX family members, auxin response transcription factors (ARFs) such as the *MP* co-orthologs *ARF4* and *ARF29*, plus candidate regulators of Kranz anatomy and vein patterning such as *TML1*, *SHR* and *SCR*. Hybridizations were carried out using transverse sections of wild-type B73 maize shoot apices, comprising the SAM (M) and the first five leaf primordia (P1 to P5) (Fig. 1A-B). These stages allowed the spatial distribution of transcripts to be captured during the development of both major (midvein and laterals) and minor (rank 1 and rank 2 intermediate) veins (Fig. 1C). Data were collected across two experiments for a total of six sections. Calcofluor White (cell wall) and DAPI (nuclear DNA) staining allowed for automatic cell segmentation with CellPose 2.0 (34) (Fig. 1B) and enabled qualitative and quantitative single-cell studies. A qualitative assessment showed consistency between Molecular Cartography and traditional *in situ* hybridizations (Figure S1). For example, *KNOTTED 1* (*KN1)* transcripts were detected primarily in the L2 layer of the meristem with very low or no expression detected in the L1 layer (Fig. S1A), consistent with published reports (35–37). Similarly, *PIN1a* (Fig. S1B) and *TML1* (Fig. S1C) transcripts were detected as previously reported, that is in all veins at all stages of leaf development (*PIN1a*) (27) or in spaces between major veins where intermediate veins are developing (*TML1*) (26). For additional validation, we also carried out a traditional *in situ* hybridization with *LAX2*, an auxin influx transporter. With Molecular Cartography, *LAX2* transcripts were detected in both major and minor veins (Fig. S1D), with higher levels detected in laterals (Fig. S1D, arrowheads) than intermediates (Fig. S1D, asterisks). These observations were consistent with results obtained by traditional *in situ* hybridization (Fig. S1E,F) (29). We thus concluded that Molecular Cartography can reliably detect transcripts in transverse sections of maize shoot apices and that the single-cell information available following cell segmentation would enable a quantitative spatial expression atlas to be generated.

### Transcript clusters distinguish functional domains in the maize shoot apical meristem

To investigate cellular relationships in the SAM, we segmented M-P1 stages and carried out qualitative and quantitative analyses of the candidate gene expression profiles within these tissues. Cluster analysis on all six sections revealed five distinct cell identities across the M-P1 stage, which includes both the SAM and P1 primordium (Fig. 2A). To spatially visualise the clusters, two representative sections were analysed: the first (E1D2) cut across the shoot apex at the base of P1 (Fig. 2B-C) and the second (S1A1) cut across the shoot apex within the region of the P0 incipient leaf and included the tip of the P1 primordium (Fig. 2B, D). Images of both sections show that one cluster corresponds to the central zone of the SAM, with high levels of *KN1* detected (yellow, Fig. 2C-D, Fig. S2). A second ‘peripheral zone’ cluster (green) is apparent either opposite a cluster corresponding to the developing P1 primordium (red, Fig. 2C) or completely encircling the central zone except for a few cells with leaf primordium identity (Fig. 2D, asterisks). Cells classified as dividing, due to the expression of genes encoding cyclins, primarily localise within the developing P1 primordium (blue, Fig. 2A-D). This observation is consistent with a rapid increase in cell number during early stages of leaf development. Finally, a handful of cells mark the boundary between the central zone of the meristem and the developing P1 primordium (magenta, Fig. 2A, C). Although the nature and function of these boundary cells in not clear, the other four clusters clearly demarcate regions of the SAM with distinct functions. Given that leaf primordia are initiated within the peripheral zone, we next sought to find traces of P0 identity in the peripheral zone opposite the tip of the P1 primordium (Fig. 2B). In *Brachypodium* and tomato, PIN1d generates the auxin maximum that determines the position of new leaves on the flanks of the SAM (38, 39). In our dataset, cells with the highest combined expression of both *PIN1d* (Fig. 2E) and *LAX5* (Fig. 2F) were positioned within the peripheral zone opposite the P1 primordium tip (Fig. 2G, Table S2). We therefore speculate that high levels of *LAX5* and *PIN1d* mark the position of P0 in the maize meristem.

**Figure 2:**
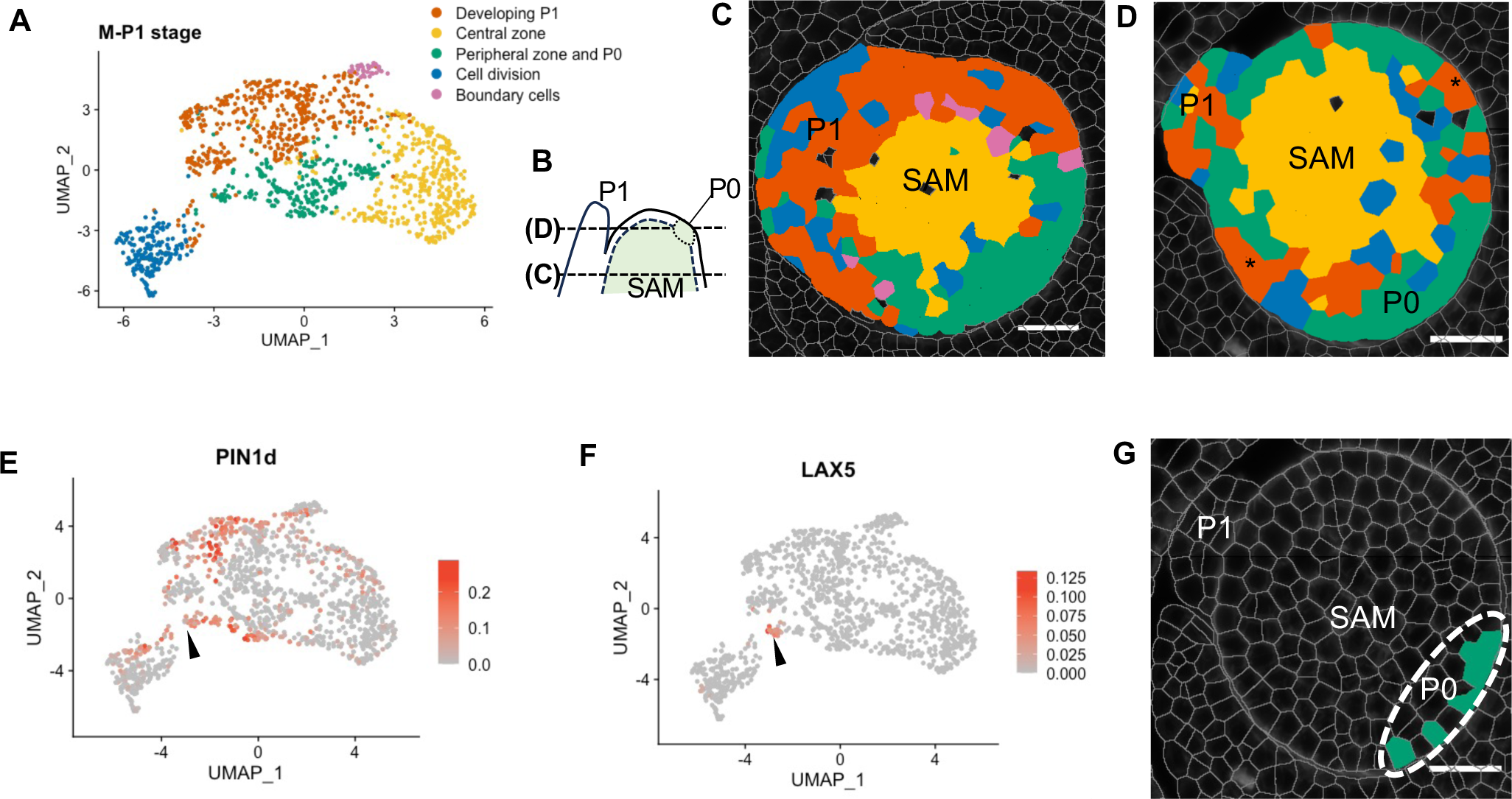
*PIN1d* and *LAX5* transcripts mark the P0 primordium. **A)** Cluster analysis of cells in the SAM and P1 primordium reveals five domains. **B)** Schematic of shoot apex showing the position of transverse sections illustrated in (C) and (D). **C)** The section taken at the base of the P1 primordium (E1D2) shows that the central zone (*KN1*-expressing) cluster localizes between the developing P1 primordium (red) and the peripheral zone (green). **D)** The section taken at the tip of the P1 primordium (S1A1) shows that the peripheral zone cluster (green) surrounds the central zone cluster (yellow), and includes the putative incipient primordium P0 region. Asterisks highlight cells with primordium identity flanking the P0 region. **E-F)** A subset (arrowhead) of cells from the ‘peripheral zone’ cluster that co-express *PIN1d* (E) and *LAX5* (F) mark the incipient P0 primordium (G, dotted ellipse). Transcript levels and cell labels are reported in Table S2. Scale bar = 30 μm.

### Leaf initiation and midvein formation are marked by distinct auxin transporters

The role of auxin transporters in directing auxin towards sites of both leaf and midvein initiation is well documented in eudicots, with PIN1 playing a dominant role. To investigate whether specific auxin transporters are associated with leaf and/or midvein initiation in maize, we compared transcript accumulation profiles across the SAM and P1. Figure 3 shows hybridization of *PIN1d*, *PIN1a*, *LAX3* and *LAX5* to transverse sections of two regions of the SAM, along with a visual summary of our observations in a deduced longitudinal view. To verify our predicted P0 site, we first examined *KN1* transcript accumulation. Importantly, transcripts were detected in the inner layers of the shoot apical meristem (Fig. 3D-F) with near absence of accumulation in our predicted P0 region (Fig. 3E, dotted line). For the genes encoding efflux transporters, *PIN1d* transcripts were detected throughout the meristem and P1 (Fig. 3G-I), with the highest signal intensity at P0 (Fig. 3H). By contrast, *PIN1a* transcripts were localised to the central zone of the meristem and were restricted to the midvein position in the P1 primordium (Fig. 3J-L, asterisks). Notably, *PIN1a* transcript accumulation appeared to be contiguous between P0 and the centre of the meristem, suggesting the presence of an auxin conduit between P0 and the vasculature of the stem. Similar to the efflux carriers, the two influx transporters also showed distinct profiles. *LAX3* (Fig. 3M-O) and *LAX5* (Fig. 3P-R), which are co-orthologues of Arabidopsis *AUX1* and *LAX1* (18), were both detected in the peripheral zone of the meristem but not in the central zone or in the P1 primordium. However, *LAX5* transcripts were detected specifically in the newly forming P0 primordium (Fig. 3Q) whereas *LAX3* was detected both within and beyond the P0 region (Fig. 3N). Collectively these results reveal that *PIN1d* and *LAX5* mark the position of the P0 primordium within the peripheral zone of the SAM whereas *PIN1a* marks both the region in the SAM where the developing midvein will connect the new leaf to the vasculature of the stem, and the region in the P1 primordium where the midvein develops.

**Figure 3:**
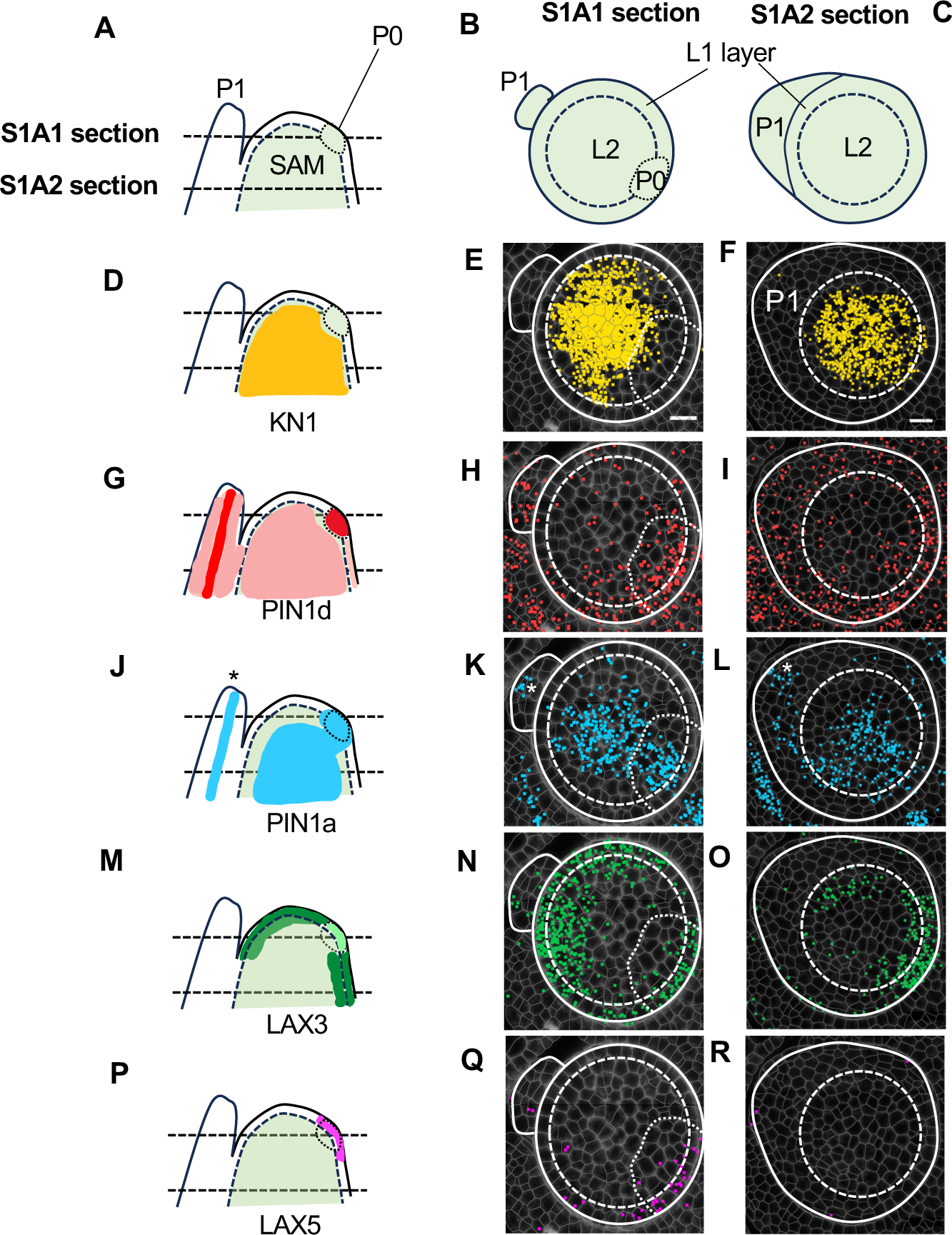
Distinct auxin transporter profiles mark leaf initiation and midvein specification. **A-C)** Schematics of sections in longitudinal (A) and transverse view (B, C). P0 = P0 primordium, P1 = P1 primordium, SAM = shoot apical meristem, L1 and L2 define the two layers within the shoot apical meristem. **D-F)** Schematic (D) and spatial transcriptomics (E, F) of *KN1*. Transcripts accumulate in the L2 layer of the shoot apical meristem, with very low or no transcript levels detected in the L1 layer or at the site of P0 initiation (dotted region, E). **G-I)** Schematic (G) and spatial transcriptomics (H, I) of *PIN1d*. Transcripts accumulate predominantly in the initiating P0 primordium (P0) but also in the SAM and the developing P1 primordium (H, I). **J-L)** Schematic (J) and spatial transcriptomics (K, L) of *PIN1a*. Transcripts are detected at the site of the developing midvein in P1 (asterisks), in the initiating P0 primordium and in the central zone of the SAM. **M-O)** Schematic (M) and spatial transcriptomics (N, O) of *LAX3*. Transcripts are detected in the peripheral zone of the SAM, including the P0 initiation site, but not in the developing P1 primordium. Lower transcript levels are detected in the P0 primordium (N) than in the rest of the peripheral zone. **P-R)** Schematic (P) and spatial transcriptomics (Q, R) of *LAX5*. Transcripts are only detected in the initiating P0 primordium, primarily in the L1 layer. Scale bars = 20 μm.

### A transcript cluster corresponding to major vein identity emerges at P2

Whereas the midvein position is determined by P1, lateral veins continue to be specified during P2 and thus to investigate cellular relationships at this later stage, we segmented P2 primordia and quantified transcript levels in each cell. Cluster analyses revealed five cellular identities, which we labelled based on the transcripts present and the spatial localisation within the primordium (Fig. 4A-C, Fig. S4). Through a visual analysis of the cluster distributions within the primordium, three cellular identities appeared to coincide with the epidermis, ground tissue and vascular tissue respectively. Dividing cells (as defined by expression of genes encoding cyclins) were present in all tissue types, although at variable levels. Intriguingly, the profile of cells in the epidermal cluster varied slightly between experiments. In one experiment there was a clear subdivision into two clusters (Fig. 4A-B, Fig. S2I-L): an abaxial epidermal cluster (grey) and an adaxial epidermal plus leaf margin cluster (cyan). The abaxial epidermis was characterized by high transcript levels of an epidermal marker (*MYB67*) and several ARFs (*ARF16, ARF34*, *ARF23*, Fig. 4D), whereas the adaxial epidermis and leaf margin cluster displayed high levels of *ANT4*, *PIN1d*, *LAX3* and *HB52* transcripts. This subdivision of the epidermal tissue was not as marked in a second experiment, where an epidermal cluster was identified but there was no clear subdivision between abaxial and adaxial/margin domains (Fig. S2G-H). Notably, differences between experiments were also observed in overall *PIN1d*, *ARF16* and *ARF34* levels (Fig. S3), suggesting that the detection of these transcripts may be more sensitive to subtle changes and variations between experiments, or to the precise cutting plane of the sections or developmental stage of the plant. In both experiments, cells in the ground tissue predominantly accumulated *ARF4* (*MP*), *ANT4* and *LAX1* transcripts, as well as low levels of *RVN1*, *SCR1/SCR1h* and *TML1* (Fig. 4D).

**Figure 4:**
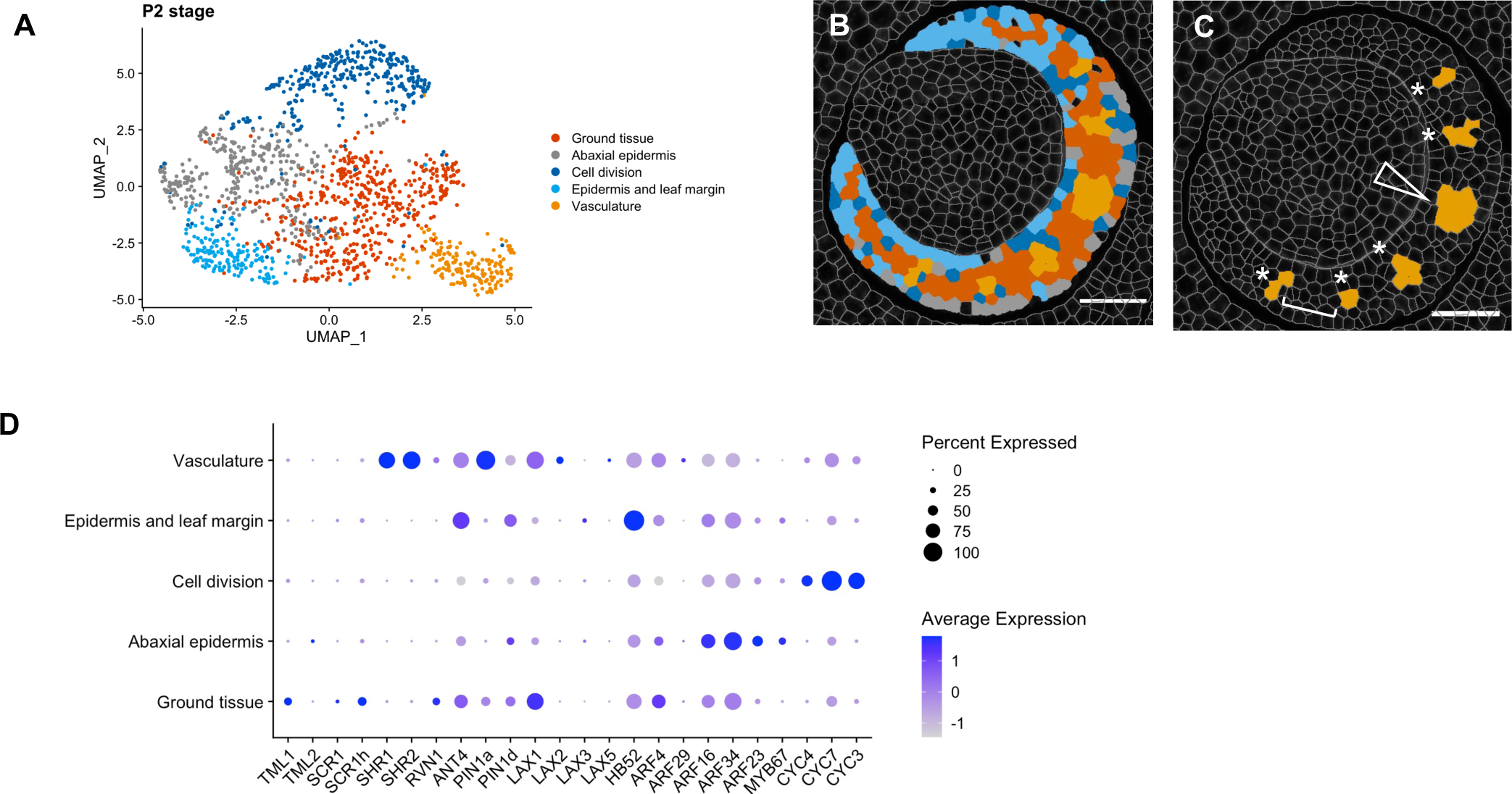
A transcript cluster with vascular identity emerges at P2. **A)** Cluster analysis and UMAP plot for the P2 leaf primordium reveals five domains. **B)** Representative image of cluster localisation in a P2 primordium, colour-coded to match (A). **C)** The vasculature cluster marks the midvein (arrowhead) and the developing lateral veins (asterisks). White bracket indicates spacing between veins. **D)** Dot-plot summarising transcript levels in each cluster. Average expression refers to the average transcript level in a cluster (cell type), calculated from the scaled data. Percent expressed refers to the percent of cells in a cluster in which transcripts (as indicated on the x-axis) were detected. Note high levels of *SHR1*, *SHR2*, *ANT4*, *PIN1a* and *LAX2* in the vasculature cluster. Scale bars = 50 μm.

Unlike the epidermis and ground tissue, which are derived from the SAM and are present at P0, procambial initial cells that give rise to the leaf veins have to be specified from within the ground tissue at successive stages of primordium development. Our data reveal that a cell cluster with vascular identity emerges at P2 (orange, Fig. 4B-C) and that it includes both the midvein (Fig. 4C, arrowhead) and lateral veins (Fig. 4C, asterisks). The cluster is characterized by high levels of *PIN1a*, *SHR1*, *SHR2*, and *ANT4* transcripts and, to a lesser extent, *LAX1* and *LAX2*. Notably, the distance between veins identified by this cluster is consistently four to five cells (Fig. 4C, bracket), which suggests that venation patterns and the distance between veins are specified by P2. Also of note is that the midvein and lateral veins form a single cluster despite representing two distinct vein ranks and developing at different timepoints (Fig. 4C). Collectively, results show that with the gene panel used in this study the P2 maize leaf primordium can be broadly subdivided into three major tissue types: ground tissue, epidermis and vasculature, with vascular identity marked by the accumulation of *PIN1a*, *SHR1*, *SHR2* and *ANT4* transcripts.

### TML1 marks a distinct intermediate vein cluster at P3 and P4

Given that all major veins can be identified by a specific transcript cluster at P2, we next investigated whether minor vein ranks can be identified by the same or a different gene set. For this purpose, cluster analysis was carried out with data from P3 and P4 primordia, to capture the development of rank-1 intermediate veins. Analysis of P3 and P4 primordia identified six and seven clusters respectively (Fig. 5A-D). In the context of the epidermis, as for P2, discrepancies were seen between experiments in terms of whether the epidermis was divided into distinct zones: one cluster associated with the abaxial epidermis and another with both the adaxial epidermis and the leaf margin (Fig. 4A,B; Fig. 5A,B,E,F; Fig S2M-R – grey and cyan respectively). In both experiments, however, the epidermal cluster was distinct from the ground tissue, which again was similar to that seen at P2 with *PIN1a*, *LAX1*, *RVN1*, *ANT4*, *HB52*, *ARF4*, *ARF16* and *ARF34* transcripts detected (Fig. 4D, Fig. 5C,D). The striking difference between clusters identified at P2 versus those identified at later stages was that two distinct vascular clusters were evident at P3 and P4. Notably these two vascular clusters were also evident when data were integrated across all stages of development (M-P1 to P5) (Fig. S5) as opposed to being analyzed within individual primordia.

**Figure 5:**
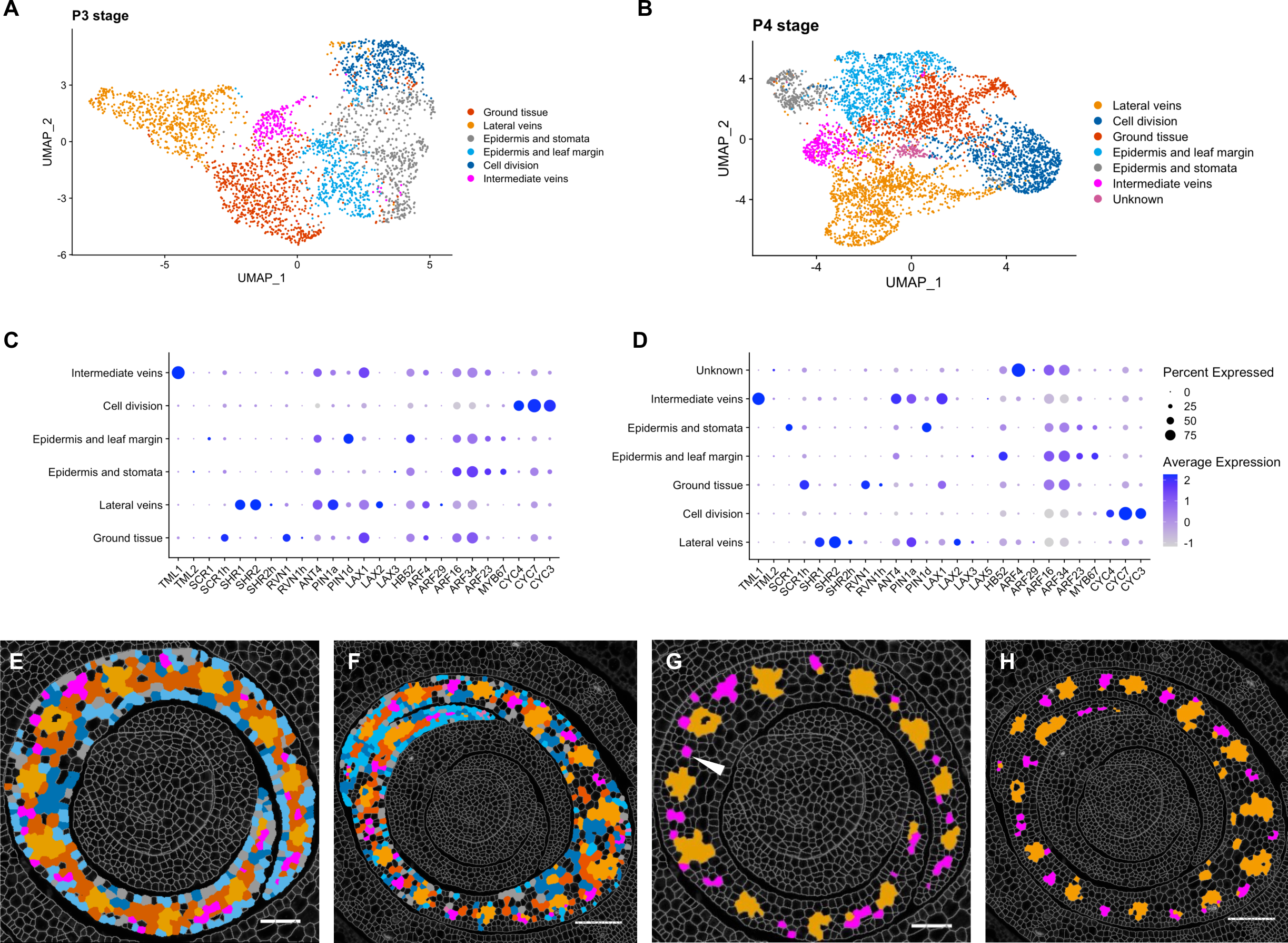
Intermediate veins cluster separately from lateral veins at P3 and P4. **A, B)** Cluster analyses and UMAP plots of P3 (A) and P4 (B) leaf primordia. **C, D)** Dotplots summarising transcript levels for each gene in each cluster at P3 (C) and P4 (D). Average expression refers to the average transcript level for each gene in a cluster (cell type), calculated from the scaled data. Percent expressed refers to the percent of cells in a cluster in which transcripts (as indicated on the x-axis) were detected. **E, F)** Representative image of cluster localisations in a P3 (E) and P4 (F) primordium, colour-coded to match (A, B). **G, H)** Two distinct vein clusters are identified, lateral (orange) and intermediate (magenta), at both P3 (G) and P4 (H). Arrowhead in G) indicates a potential procambial initial. Scale bars = 50 μm.

The spatial localisation of the two vascular clusters and the spacing between them within the P3 and P4 primordia suggest that they represent distinct lateral (Fig. 5G,H - orange) and intermediate (Fig. 5G,H - magenta) vein identities. Both clusters share a common set of transcripts which include *PIN1a*, *LAX1*, *ANT4*, *PIN1d*, *HB52*, *ARF16* and *ARF24*. However, the intermediate vein cluster is distinguished by high levels of *TML1* transcript (Fig. 5C,D), which is consistent with the observation that the *tml1* mutation in maize specifically perturbs the development of rank-1 intermediate veins (Fig. S1C) (26). The intermediate vein cluster also displays lower levels of *SHR1* and *SHR*2 transcripts than the lateral vein cluster (Fig. 5C,D). In conclusion, clustering of cells from stage P3 and P4 highlighted similarities and differences in the development of different vein ranks. Both lateral and rank-1 intermediate veins accumulate a similar set of auxin transporters and procambial markers, such as *PIN1a*, *LAX1*, *ANT4* and *HB52*. However, the clear difference between the lateral and rank-1 intermediate vein clusters, and the specific accumulation of *TML1* transcripts in the intermediate cluster, suggest that vein ranks are transcriptionally distinct after just a few divisions of the procambial initial.

### Vein formation across the medio-lateral leaf axis reveals a temporal hierarchy of PIN1a, SHR1 and LAX2 accumulation during early procambium development

To determine the extent to which the spatial regulation of genes associated with vein formation is accompanied by temporal regulation, we analysed the distribution of transcripts across the shoot apex. For this analysis we focused on genes that are shared between lateral and intermediate veins clusters, such as *PIN1a*, *SHR1*, *SHR2* and *ANT4,* in order to dissect events during the first few divisions of the procambial initial. We discovered that *PIN1a* was the earliest and most persistent marker of vein development, being detected at all stages of development from M-P1 to P5 (Fig. 6A). In M-P1, *PIN1a* transcripts marked both the position of the developing midvein (Fig. 6B, white ellipse, “m”) and the central zone of the meristem, suggesting a role in connecting the newly developing midvein with the vasculature of the stem. From P2 onwards, transcripts were detected in all vein ranks of all leaf primordia, i.e. the midvein (Fig. 6C-D, white ellipse), laterals (Fig. 6D, empty arrowheads) and intermediates (Fig. 6D, filled arrowheads). Notably, *PIN1a* transcripts were detected before and after procambial centres became anatomically distinguishable from the surrounding ground tissue, with other procambial and vascular markers following the appearance of *PIN1a*. For example, we detected transcripts of the GRAS transcription factor *SHR1* (Fig. 6E, Fig. S6A-E) in all vein ranks (as indicated by *PIN1a* expression), but at a slightly later stage of development. These observations revealed not only that the expression of *SHR1* occurs temporally after that of *PIN1a*, but also that the development of veins across the medio-lateral axis follows a specific dynamic: veins closer to the midrib express *SHR1* earlier than those closer to the leaf margin. In a similar manner, we observed that *LAX2* transcripts accumulated at least one plastochron later than *SHR1* (Fig. 6F, Fig. S7A-E). By contrast, however, *ANT4* is present in all intermediate veins across the medio-lateral axis of the leaf (arrowheads, Figs. S6F, S7F). This suggests that the observed spatio-temporal dynamic is specific for *SHR1* and *LAX2*. Together, our data suggest that PIN1a is the earliest leaf procambial marker at the developmental stages considered in this study and that other procambial markers follow in a specific temporal and spatial hierarchy across the medio-lateral axis.

**Figure 6:**
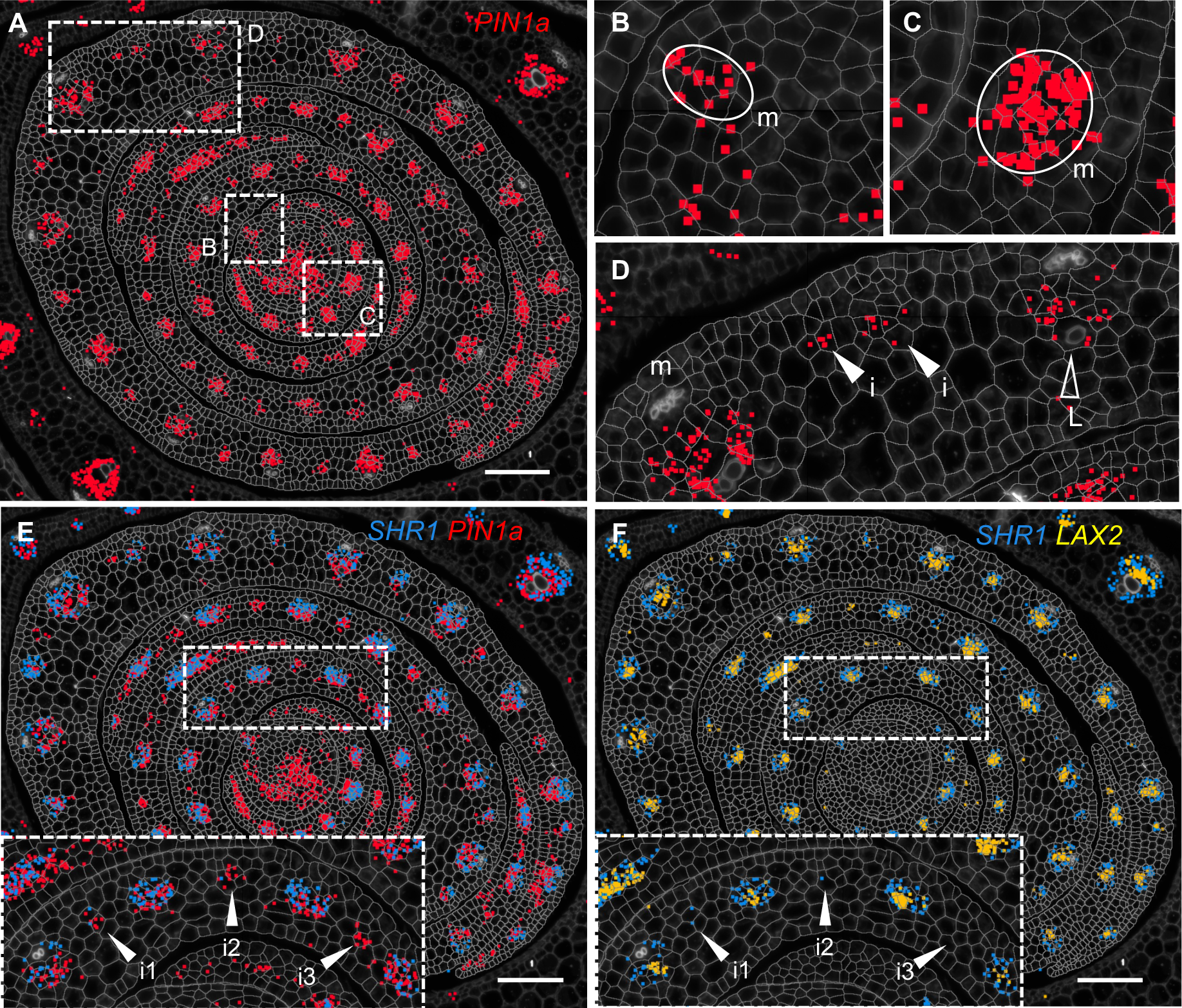
*PIN1a* accumulation precedes the sequential accumulation of *SHR1* and *LAX2* transcripts in dividing procambial centres. **A)** Representative image of *PIN1a* transcript accumulation (red pixels) in the SAM and P1 to P5 leaf primordia. Transcripts accumulate in both the leaf sheath and leaf blade, at all stages of vein development and in all vein ranks. White boxes represent the position of enlargements shown in (B-D). **B, C)** *PIN1a* is detected at low levels in the future midvein at P1 (B - m, circled) and at higher levels at P2 (C-m, circled). **D)** *PIN1a* transcripts are detected in all developing veins within the P5 primordium (m, midvein; i, intermediate; L, lateral). **E)** Superimposed images of *PIN1a* (red) and *SHR1* (blue) transcripts reveal that *PIN1a* accumulation precedes that of *SHR1*. Arrowheads in the inset indicate three developing intermediate veins (i1, i2, i3), in order of their initiation across the medio-lateral leaf axis. *SHR1* transcripts are detected in i1 and i2, but not in the more recently developed i3. **F)** Superimposed images of *SHR1* (blue) and *LAX2* (yellow) transcripts reveal that *SHR1* accumulation precedes that of *LAX2*. Arrowheads in the inset indicate the same developing intermediate veins as in (E): *LAX2* transcripts are not detected in any of these veins but are found in neighbouring older veins. Scale bars = 100 μm.

### Vascular markers become polarized in the adaxial-abaxial axis at late stages of vein development

Having observed that at least a subset of venation markers are sequentially expressed during procambium initiation, we next investigated whether localisation could also vary between early and late stages of vein development. We took into consideration two vein ranks, the midvein and lateral veins, for which a trajectory from early specification (M-P1 and P2 stage) to phloem and xylem differentiation (P3 to P5) could be traced across the developmental stages in our study. Several markers for which transcripts are localised in all vein ranks were analyzed. A representative example is provided in Figure 7A, where *PIN1a* (red), *HB52* (yellow) and *LAX2* (blue) transcripts are superimposed, and all transcripts are detected in all vein ranks and at all leaf developmental stages. Regardless of vein rank, at stages prior to differentiation of phloem or xylem, all three markers were detected throughout the procambial centre (Fig. 7B). Once phloem and xylem were differentiated, however, *PIN1a*, *HB52* and *LAX2* transcripts were polarly localized (Fig. 7C,E). In both the mid- (Fig. 7C) and lateral (Fig. 7E) veins, *PIN1a* transcripts were detected on the adaxial side, *HB52* transcripts towards the phloem on the abaxial side, and *LAX2* juxtaposed between the two. Collectively, our data suggest that vascular-associated genes are tightly regulated temporally and spatially, both within and between different vein ranks, with changes in spatial localisation marking transitions between the specification of procambial initial cells, the development and extension of procambial strands, and differentiation of tissues within the vein.

**Figure 7:**
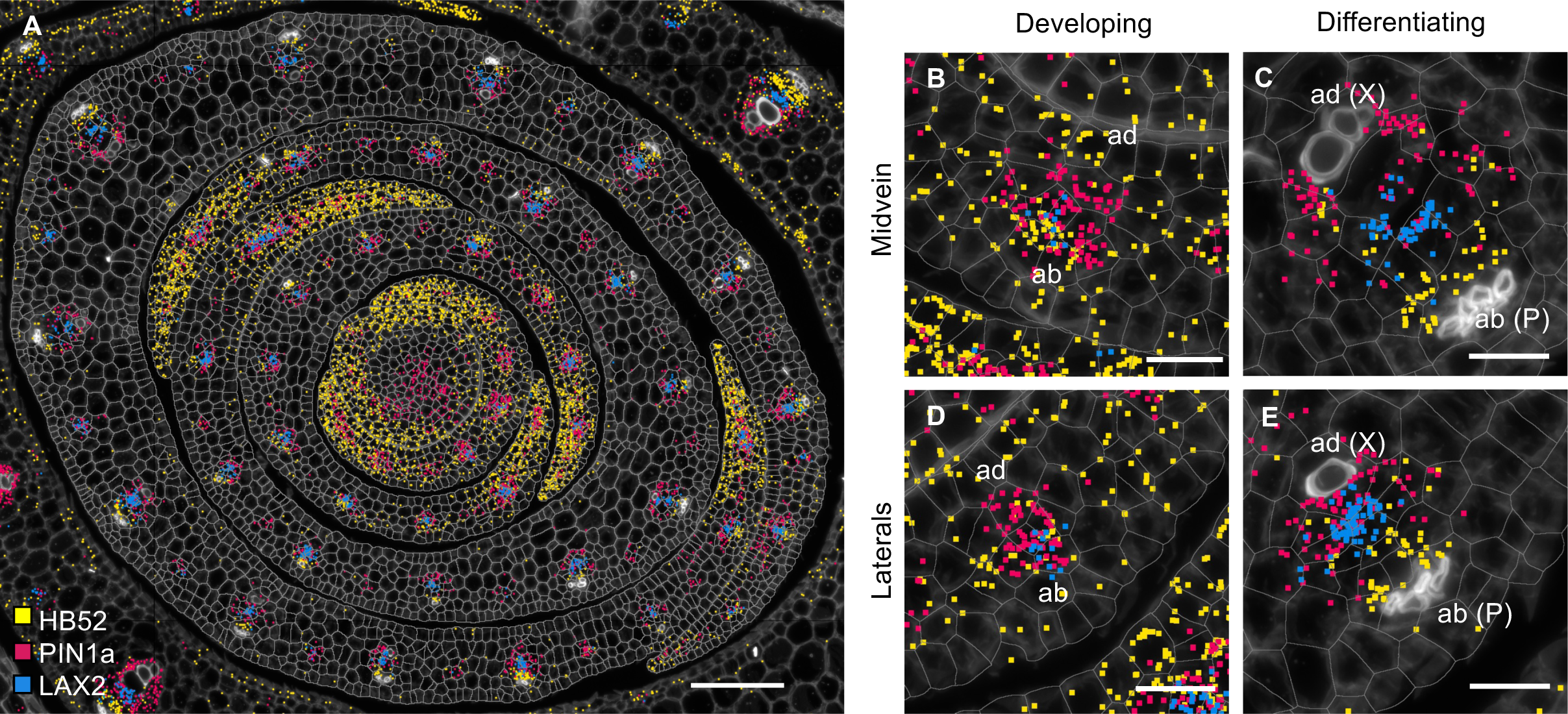
Differentiation of phloem and xylem is accompanied by polar localization of vascular markers. **A-E)** Transcripts of vascular markers *HB52* (yellow), *LAX2* (blue) and *PIN1a* (red) accumulate in developed and developing veins in all leaf primordia (A). All markers co-localise in developing mid-(B) and lateral (D) veins, with very little adaxial (ad) or abaxial (ab) separation. In differentiated mid-(C) and lateral (E) veins, transcripts preferentially accumulate on the phloem-side (P, as in the case of *HB52*, yellow), xylem-side (X, as in the case of *PIN1a*, red), or juxtaposed (as in the case of *LAX2*, blue). Scale bars = 100 μm (A), 20 μm (B-E).

## DISCUSSION

Here we have used multiplexed *in situ* hybridization and a set of marker genes associated with auxin transport and vein patterning to investigate cellular relationships during leaf specification and vein formation in maize shoot apices (Fig. 1). On leaf initation, we reveal that it is possible to identify the region corresponding to incipient P0 leaves within the peripheral zone of the SAM, based on high density of *PIN1d* and *LAX5* transcripts (Fig. 2). *PIN1a* transcripts connect the site of the incipient leaf with the central zone of the meristem, likely connecting the new leaf with the vasculature of the stem and determining the position of the midvein at P1 (Fig. 3). From P2 to P4, the specification of both major (midvein and laterals) and minor (intermediate) vein ranks can be traced, with intermediate veins characterized by the presence of the transcription factor *TML1* (Fig. 4, 5). Strikingly, the localisation of the procambial marker *PIN1a* and the spatial position of major and minor vein clusters suggest that vein spacing could be pre-patterned prior to *PIN1a* accumulation. Once *PIN1a* accumulation (and presumably auxin flux along the proximo-distal leaf axis) is initiated, a spatiotemporal hierarchy is triggered in which vein markers accumulate in a sequential manner that is position-dependent across the medio-lateral leaf axis (Fig. 6). During the final stages of vein development, in which phloem and xylem are differentiated, vascular markers become spatially restricted in the adaxial-abaxial leaf axis (Fig. 7). Collectively these data suggest progressive restriction of gene expression in all three leaf axes as veins are specified, initiated, extended and then differentiated.

The importance of understanding cell lineage relationships during development has long been recognised, not least to distinguish mechanisms in which cell fate is inherently specified in descendents following mitosis, from those in which fate specification relies on positional information from neighbouring cells (reviewed in (40)). In maize, many aspects of leaf development have been elucidated by exploiting genetic mosaics, including the presence of distinct central and marginal lineages in the medio-lateral leaf axis (41, 42) and the presence of a lineage in the centre of the adaxial-abaxial axis from which all veins are derived (7). With the advent of single cell sequencing, spatial transcriptomics and quantitative *in situ* hybridizations, discussion of lineage relationships have altered. Cells with shared fates are identified regardless of whether that fate is determined by lineage-based or positional mechanisms (33, 43–45). Importantly, combining information from both approaches allows classical lineage distinctions to be associated with specific gene expression profiles and also facilitates more refinement of cellular relationships during later stages of development. For example, we identified transcriptionally distinct populations of cells in the peripheral zone of the SAM that will give rise to the P0 primordium, and identified two distinct populations of vein-forming procambial cells in the central ground meristem layer of P3 and P4 leaf primordia. Notably, some of the markers that defined developing procambial lineages, such as *PIN1a*, *TML1* and *HB52*, could be detected in a dispersed pattern prior to vein specification. For example, *TML1* transcripts were detected at P2, prior to intermediate vein specification, but accumulation patterns were not restricted to the central ground meristem layer (Fig. S1C) (26). Similarly, *TML1* transcripts were detected primarily in procambial cells that were dividing to form intermediate veins in the middle of the P3 primordia (arrowheads, Fig. S1C), but were detected in a more dispersed pattern at the leaf margins where procambium was still being specified. Undoubtedly, better refinement of the developmental trajectories of cell lineages within and between leaf primordia could be achieved using unbiased approaches such as scRNAseq (44, 46) and/or 3D analyses (31) that would avoid variability between sections. However, even this limited gene set has enabled the identification of transcriptionally distinct cell populations within the vein-forming cell lineage (*sensu stricto*) in the middle of the adaxial-abaxial leaf axis (7) and has provided information about when cell fate is specified.

In the context of vein patterning, cellular relationships identified through transcript profiles have revealed distinctions that have not been made previously by either cell lineage or histological analyses. The set of genes used in this study distinguished two separate cell clusters between stages P2 and P4, corresponding to major (midvein and lateral) and intermediate veins (Figs. 4, 5). No substantial differences were detected between midvein and lateral vein clusters, which were grouped together from P2 onwards. This observation could reflect the fact that our dataset does not include genes which are specific for either midvein or lateral vein identity. Alternatively, it suggests that the midvein and lateral veins are not intrinsically different, a suggestion supported by the phenotype of midribless mutants (47, 48). By contrast, minor veins are transcriptionally distinct from major veins once procambium is specified, as evidenced by transcript localisation profiles (see example of single cells in Figure 5G) and the phenotype of loss of function *TML1* mutants (26). Although both minor and major veins are specified *de novo* from the middle layer of the ground meristem, this observation suggests that procambial initials are inherently distinct. This hypothesis is supported by the observation that intermediate veins start to be specified near the midvein during P3, when growth in the medio-lateral axis is ongoing and lateral veins are still being specified near the leaf margin (Fig. 5 G-H, orange). This leads to a gradient in the medio-lateral axis, with veins closer to the midrib being more developed than those closer to the leaf margin (11). If major and minor vein lineages were not inherently distinct, intermediate veins close to the midrib would be of similar size to lateral veins close to the leaf margin but they are not (Table S3). These findings suggest that procambial initial cells that give rise to major and minor vein ranks in the maize leaf are distinct at inception.

In the model eudicot *Arabidopsis thaliana,* auxin fluxes have been associated with vein patterning, with pre-existing vasculature used as a positional landmark towards which auxin flows and new procambium is initiated (12, 17). In contrast, the localisation of transcripts encoding auxin transporters in maize suggests that auxin fluxes associated with procambial initiation are pre-patterned. Specifically, *PIN1a* transcripts are first detected in individual procambial initial cells that are already spaced appropriately across the medio-lateral axis. As such, *PIN1a* activity is not necessary to determine which cells in the ground tissue will commit to procambial fate. Notably, the DR5 transcriptional reporter is detected along the leaf margin in maize leaves, indicating that an auxin maximum is present at the margin (27, 49). Given that PIN1a does not overlap with the DR5 signal at the margin of P0 to P5 primordia (27), auxin flux from the margin is unlikely to be inducing procambial initial cell specification or procambial strand extension. Instead, an unidentified factor must be inducing PIN1a accumulation in the subset of regularly spaced ground tissue cells that become procambial initial cells. Once *PIN1a* is activated, other auxin transporters and vascular markers are detected in developing veins but temporal and/or spatial patterns of transcript accumulation again suggest progressive restriction of activity. For example, maize abaxial and adaxial markers *ARF3a* (*AUXIN RESPONSE FACTOR 3a*) and *RLD1* (*ROLLED LEAF1*) are localised throughout the adaxial (RLD1) or abaxial (ARF3a) leaf domains between P0 and P3, but by P4 they are primarily restricted to the vasculature and are associated with differentiated xylem (*RLD1*) and phloem (*ARF3a*) (49). Together, these findings suggest that the position of procambial initials in the medio-lateral axis of the maize leaf is pre-patterned prior to auxin flux through the procambial strand, and that the patterning of differentiated tissues within each vein is similarly pre-patterned in the adaxial-abaxial leaf axis.

The role of auxin in both leaf initation and vein patterning has been acknowledged for some time (1, 50) but the spatial transcriptomes reported here reveal new insights. In *Arabidopsis thaliana*, *PIN1* acts as a master regulator of both leaf positioning and vein specification (12, 51, 52). In non-Brassicaceae, however, the expansion of the PIN1 family and the emergence of a sister clade (SoPIN1/PIN1d) enabled sub-functionalisation (18, 38, 39, 53). Our analysis reveals that positioning of the P0 primordium in maize is associated with combined accumulation of transcripts encoding the auxin efflux and influx carriers *PIN1d* and *LAX5* (Fig. 3), whereas the specification of procambium is associated with the accumulation of *PIN1a*, *PIN1d* and *LAX1*, with *LAX2* likely being the dominant influx carrier later in vein development. Our results are consistent with previous findings where combined expression of *PIN1a*, *PIN1d* and *LAX1* (note *LAX2* in (19)) was detected from procambial initial cell selection through to later stages of procambium development (19). Detection of *PIN1d* but not *PIN1a* at the site of the P0 primordium suggests a functional subdivision within the maize PIN1 family as seen in tomato and *Brachypodium* (38, 52), with *PIN1d* (*SoPIN*) regulating phyllotaxy and *PIN1a* tracing the midvein and its connection to the subtending vasculature (Fig. 3) (49). We further speculate that *LAX5*, co-ortholog with *LAX3* of *Arabidopsis AUX1/LAX1*, supports the role of *PIN1d* in generating an auxin maximum in the L1 of the SAM, whereas *LAX2* acts with *PIN1a* to extend procambial strands in the proximo-distal leaf axis. Together, these findings suggest a subdivision of roles within both the *PIN1* and *LAX* gene families that allows different combinations of auxin influx and efflux transporters to be recruited in different developmental contexts.

## CONCLUSIONS

This study used a quantitative multiplex *in situ* hybridization platform to identify populations of cells with shared identity, and to elucidate how those populations diverged during the course of leaf primordium development in maize. Regardless of whether they arise as the result of positional or lineage-based cues, cell fates can be associated with the spatial co-localisation of transcripts within a tissue. As such, we propose that major and minor vein ranks in the maize leaf have distinct identities at inception from the ground meristem. We further propose that veins in the maize leaf are pre-patterned in all three leaf axes, with auxin triggering downstream specification events through finely tuned spatio-temporal accumulation of influx and efflux transporters.

## METHODS

Twenty seven genes (Table S1) were selected as probes to hybridize to maize shoot apices comprising the shoot apical meristem and the first five leaf primordia. Multiplexed *in situ* hybridizatiion was carried out using Resolve Biosciences Molecular Cartography technology, and data analyses and plots were carried out with R Studio and the Molecular Cartography ImageJ plugin Polylux. See Supplementary Information for full details of Methods.

## DATA AVAILABILITY

All raw data, Calcofluor White and DAPI images and cell segmentations used in this study and in Vlad *et al* (26) are deposited on Zenodo (URL: https://doi.org/10.5281/zenodo.10605855). All unprocessed and normalised and scaled data are available in Data S1, S3 and S4, and cell identities following cluster analyses are reported in Data S2.

## AUTHOR CONTRIBUTIONS

CP and JAL conceived and designed the experiments, MZ and CP wrote the code and performed the cluster analysis, OS carried out the traditional *in situ* hybridization experiments, SB carried out the multiplexed *in situ* hybridizations under the supervision of CK, CP performed all the sample preparation and data analysis; CP and JAL wrote the first draft of the manuscript and all authors contributed to the final version.

## ACKNOWLEDGEMENTS

The authors thank all members of the Langdale lab for constructive discussions of the results and data analysis, and feedback on the manuscript. We also thank Thomas Hughes for helping with the sample preparation, and Scott Poethig, Tina Schreier, Hilde Nelissen, Jessica Joossens and Denia Herwegh for feedback on the manuscript.

