## Supplementary Information for "Multiplexed *in situ* hybridization reveals distinct lineage identities for major and minor vein initiation during maize leaf development"

### Materials and Methods

#### *Molecular Cartography probe design*

Probes were designed using Resolve BioSciences' proprietary design algorithm and gene annotations from the *Zea Mays* RefGen V4. The final set of probes was selected as outlined in (1).

#### *Plant growth and tissue preparation for Molecular Cartography*

Samples were prepared, hybridized and imaged as in Vlad et al. (2). In total six sections were analysed - E1B2, E1D2, S1B1, S1B2, S1A1 and S1A2. Two independent experiments were carried out – the first with sections E1B2 and E1D2 and the second with samples S1A1, S1A2, S1B1 and S1B2.

#### *Data analysis*

All data analyses and plots were carried out with R Studio and the Molecular Cartography ImageJ plugin Polylux proprietary of Resolve Biosciences. Unprocessed single-cell data was extracted from the spatial images using the ImageJ plugin Polylux from Resolve BioSciences. Correlation plots were generated using *corrplot* (3) to evaluate the linear relationships between genes across experimental repeats (note that each experimental repeat was also a biological repeat as each section was taken from a different shoot apex). The R package Seurat (4) was used to carry out all clustering data analyses and plotting. The Seurat clustering workflow was adapted with minor adjustments to fit the data frame from molecular cartography. Briefly, for each stage, a Seurat object was generated from unprocessed data with parameters: min.cells = 20 (P2) or 30 (M-P1, P3, P4, M to P5); min.features=3. Parameters were adjusted to account for the difference in the number of cells per primordia stage. Data was normalised using the "LogNormalize" method and a scale factor of 1. Scaling using ScaleData, to prevent overrepresentation of frequent signals in dimensional reductionwas then done using default parameters. A principal component analysis (PCA) was performed using standard parameters. After determining dimensionality of the dataset via the Seurat ElbowPlot function, 10 components were taken into consideration to run the Uniform Manifold Approximation and Projection (UMAP) dimensionality reduction to cluster the cells Clustering was based on neighbours, found using the "FindNeighbours" command (dims = 1:10, dims = 1:13 for M to P5), followed by a default "FindClusters". FindClusters allowed cluster generation by constructing a Shared Nearest Neighbour (SNN) graph and applying the Louvain algorithm. Cluster resolution was set at either 0.3 (P4, M to P5) or 0.4 (M-P1, P2, P3). Dimension plots were generated using the "DimPlot" command and "umap" as the reduction method. Cluster location was visually checked using the Polylux plugin, and identity was assigned based on the cluster localisation within the sections.

#### *Tissue fixation and preparation for traditional in situ hybridization*

Traditional *in situ* hybridization was carried out using wax-embedded wild-type B73 maize shoot apices as previously described (5) with a 268 bp digoxygenin (DIG)-RNA labelled probe complementary to the 3'UTR sequence of the *LAX2* gene (starting at 126 bp downstream of the stop codon).

#### *Vein width quantification*

Wild-type B73 maize seeds (n = 5) were germinated and grown as for Molecular Cartography, using a 3:1 mixture of John Innes #3 and vermiculite. Following full blade expansion of leaves three and four, 5 mm segments spanning the whole leaf width were harvested from the widest point of the blade. Samples were then fixed in 3:1 EtOH/acetic acid for 30 min at room temperature, followed by a 30 min wash in 70% EtOH. After an overnight incubation in 70% EtOH at 4°C, leaf segments were embedded in blocks of 6% agar and sectioned using a vibratome. Leaf sections were imaged using a stereomicroscope and the width of lateral and intermediate veins outside of the midrib region were measured using ImageJ. Lateral veins were labelled with increasing numbers across the medio-lateral axis (L1 indicating the lateral vein closest to the midrib). Average intermediate veins number and width were measured between successive lateral veins, and between the last lateral vein and the leaf margin. All vein width quantifications were carried out on one half of the leaf segments.

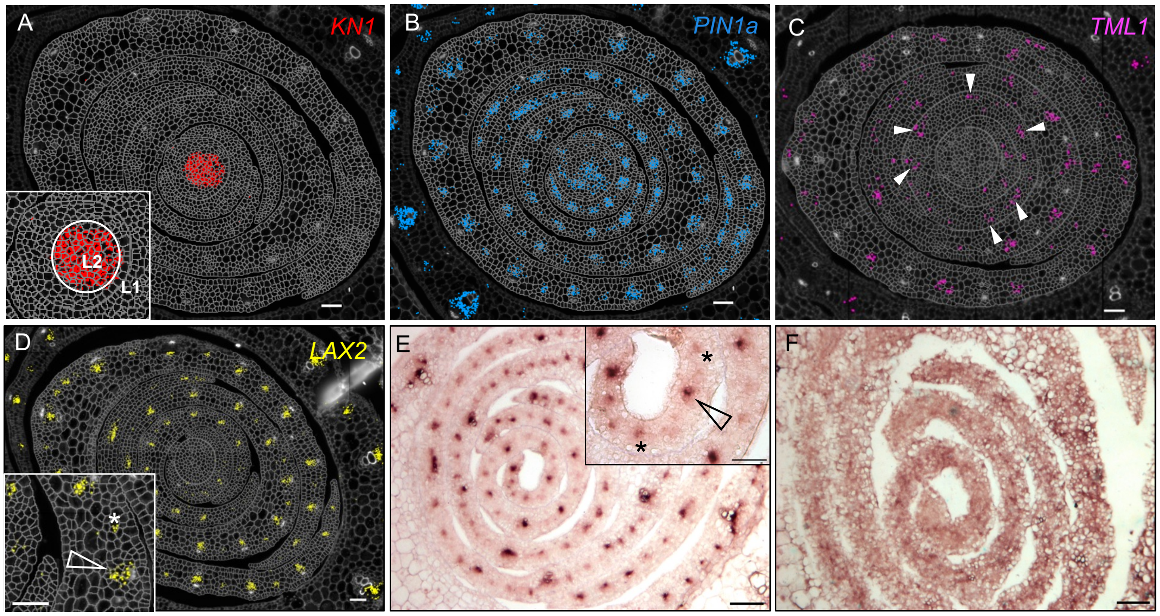


**Fig. S1. Validation of Molecular Cartography methodology for localization of transcripts in maize shoot apices**. **A)** *KN1* transcripts localize in the shoot apical meristem, specifically the L2 layer as previously shown by in situ hybridization (6). **B)** *PIN1a* transcripts mark all developing veins at all stages of leaf development. **C)** *TML1* transcripts accumulation in developing intermediate veins from stage P3 (white arrowheads). **D, E)** Molecular Cartography (D) and *in situ* hybridization (E) of *LAX2*. In both cases, transcripts are detected in developing vascular tissue with higher levels detected in major veins (inset, arrowhead) compared to minor veins (inset, asterisks). **F)** *In situ* hybridization control with sense *LAX2* probe. Scale bars = 50 μm.

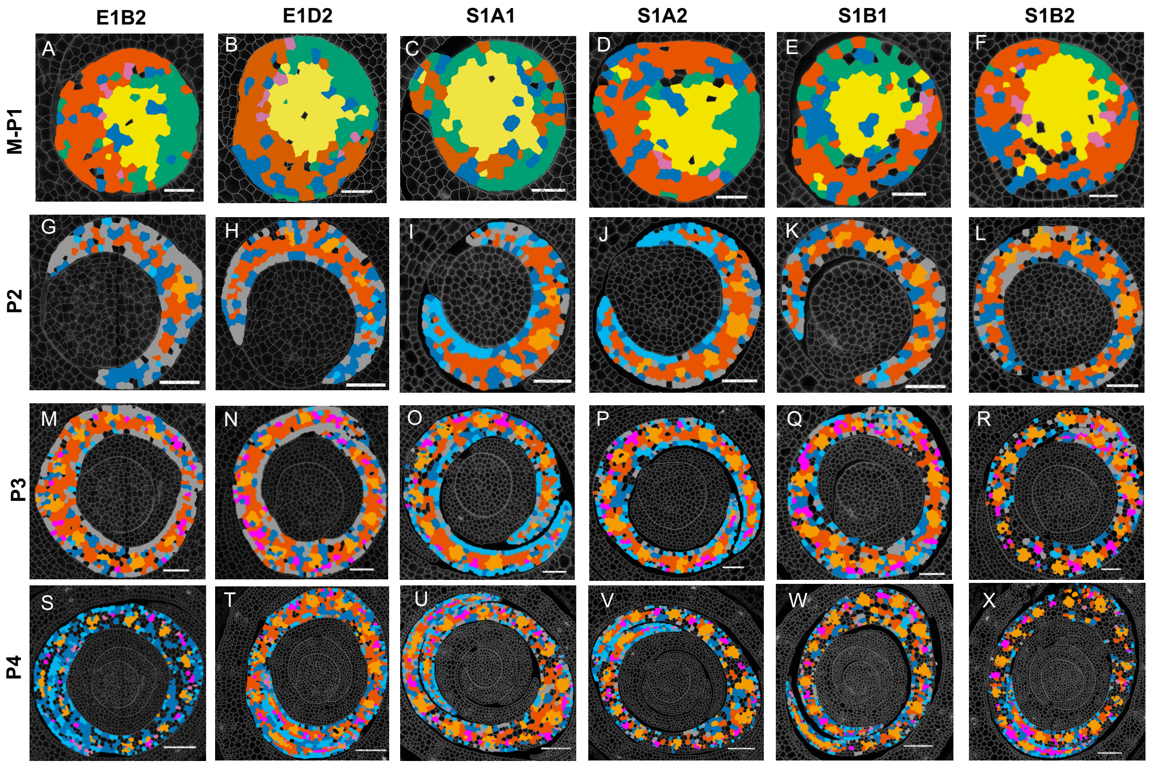


**Fig. S2. Visual representation of clusters at all stages of leaf development.** Sample numbers are depicted across the top and plastochron stages at the left. Cluster colours are as depicted in the main figures. Scale bars = 30 µm (A-F), 50 µm (G-R), 100 µm (S-X).

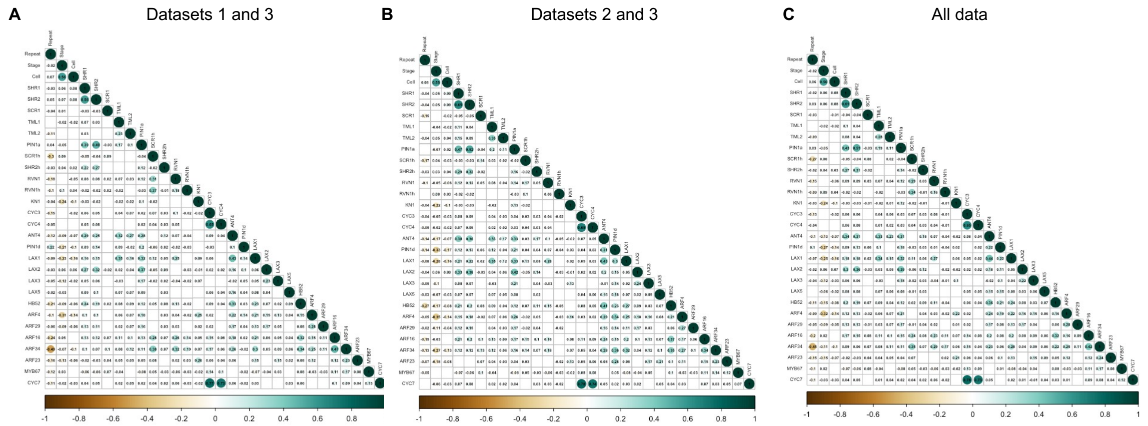


**Fig. S3. Correlation analysis of the Molecular Cartography datasets. A)** Correlation plot for comparison of different experimental datasets with the same tissue preparation method (dataset 1: E1B2, E1D2; dataset 3: S1B1, S1B). **B)** Correlation plot for comparison of datasets with manual (dataset 32) and automatic (dataset 2: S1A1, S1A2) tissue preparation method within the same experiment. **C)** All datasets together. Brown colour indicates negative correlations between rows and columns, blue indicates a positive correlation. The number in each cell represents the correlation coefficient (confidence level = 0.95).

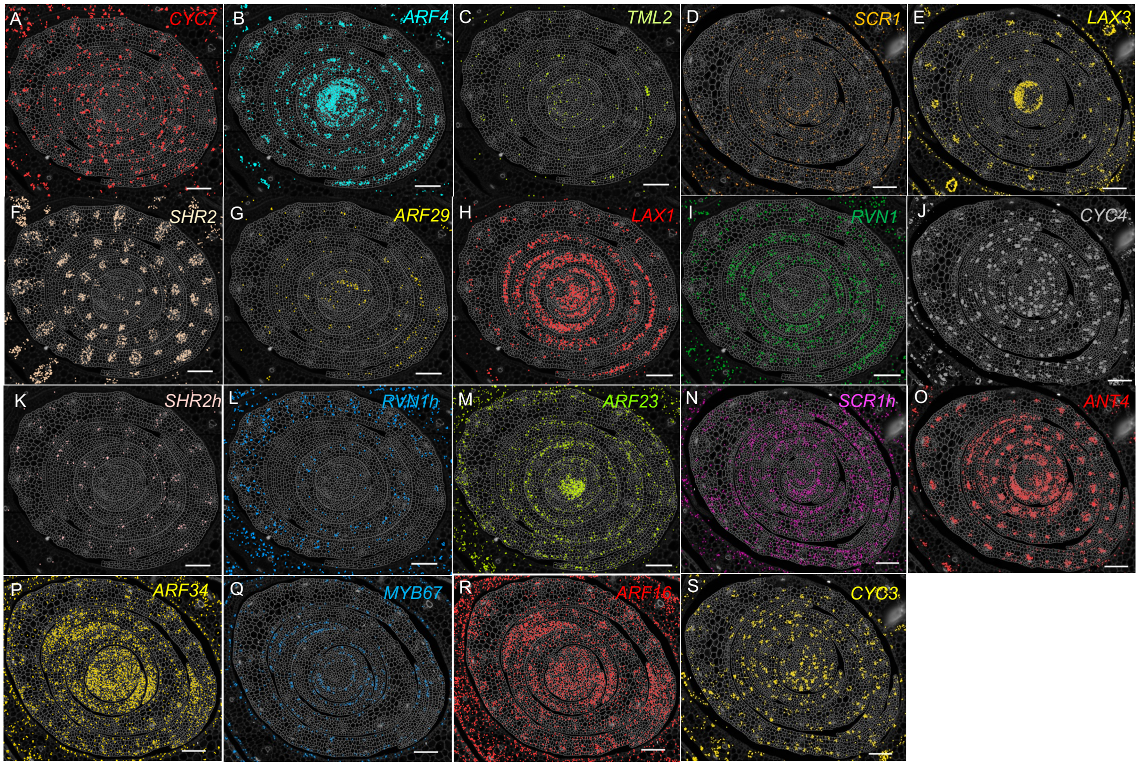


**Fig. S4. Representative images showing the localization of transcripts for all genes in the Molecular Cartography panel.** Images are chosen across all sections. Representative transcript localization for *PIN1a*, *KN1*, *LAX2*, *SHR1*, *TML1* and *HB52* are provided in Fig. S1, Fig. 6 and Fig. 7. Scale bars = 100 μm.

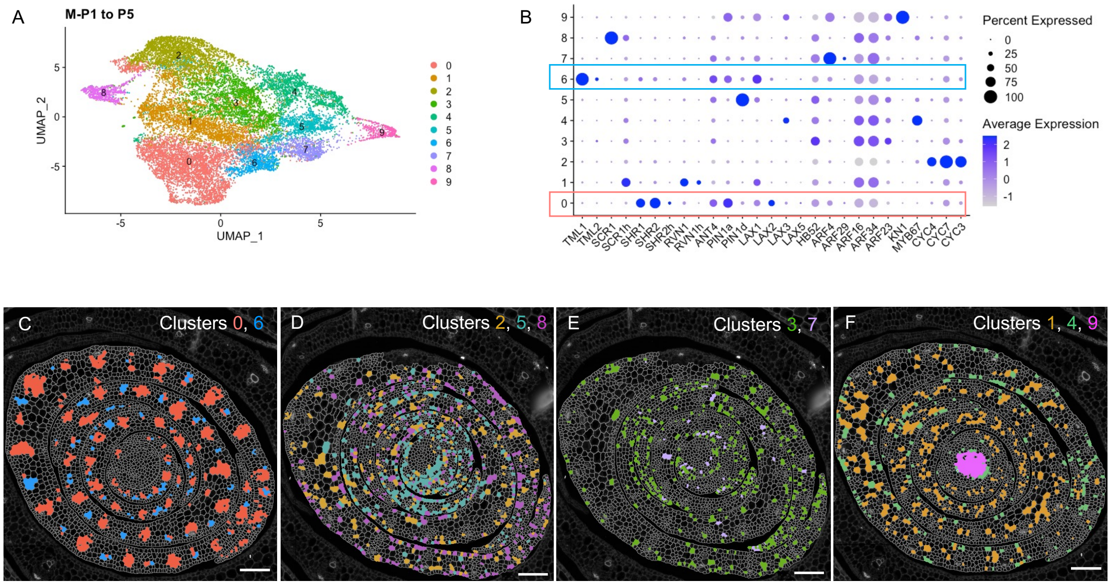


**Fig. S5. Data clustering from stage M-P1 to P5 indicates separate identities for major and minor veins. A)** Cluster analysis and UMAP plot of data collectively analysed for stage M-P1 to P5. **B)** Dotplot summarising transcript levels for each gene in each cluster at M-P1 to P5. Average expression refers to the average transcription level for each gene in a cluster (cell type), calculated from the scaled data. Percent expressed refers to the percent of cells in a cluster in which transcripts (as indicated on the x-axis) were detected. Blue and red boxes highlight the expression profiles of clusters 6 and 0. **C-F)** Representative images of cluster localisation from M to P5, colour-code as in panel A. Scale bars = 100 µm.

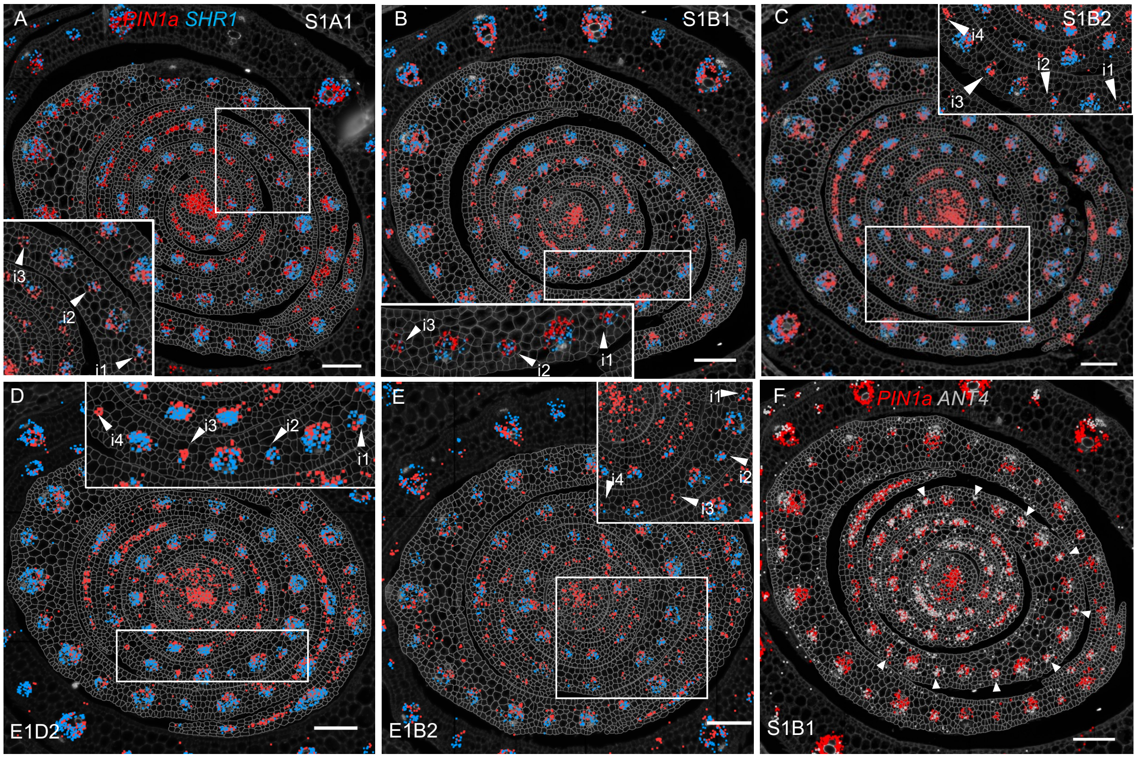


**Fig. S6. *PIN1a* expression precedes that of *SHR1* across the medio-lateral axis**. ***A-E)*** Transcript accumulation for *PIN1a* and *SHR1* is detected in both the sheath and blade and in both major and minor veins. Intermediate veins closer to the midrib accumulate *SHR1* before those closer to the leaf margin. In insets, intermediate veins are labelled with progressive numbers indicating the degree of proximity to the midrib. **F)** Unlike *SHR1*, *ANT4* is detected in all intermediate veins across the medio-lateral axis (white arrowheads). Scale bars = 100 μm.

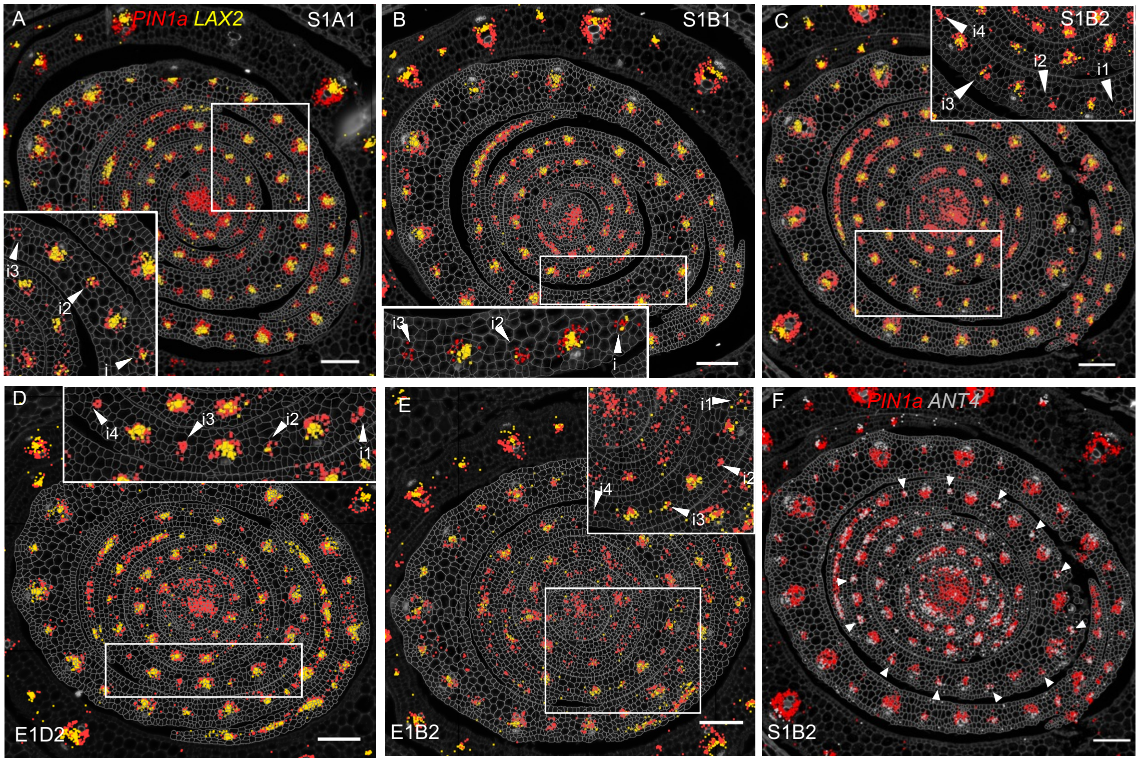


**Fig. S7. *PIN1a* expression precedes that of *LAX2* across the medio-lateral axis. A-E)** Transcript accumulation for *PIN1a* and *LAX2* is detected in both the sheath and blade and in both major and minor veins. Intermediate veins closer to the midrib accumulate *LAX2* before those closer to the leaf margin. In insets, intermediate veins are labelled with progressive numbers indicating the degree of proximity to the midrib. **F)** Unlike *LAX2*, *ANT4* is detected in all intermediate veins across the medio-lateral axis (white arrowheads). Scale bars = 100 µm.

**Table S1. List of genes used for Molecular Cartography.**

| Gene_name | Gene_name Maize GDB | Gene ID (v3) | Gene ID (v4) | Gene ID (v5) | Origin of gene name (*supported by phylogeny) | Orthology | Variations |
|---|---|---|---|---|---|---|---|
| *TML1* | dot5 | GRMZMG2150011 | Zm00001d020037_T001 | Zm00001eb309530 | Vlad et al 2024* | AtWIP6 | Previously refrred to as Zmdot1 or Zmdot5 |
| *TML2* | – | GRMZM2G134998 | Zm00001d005566_T001 | Zm00001eb097850 | Vlad et al 2024* | AtWIP6 | Previously refrred to as Zmdot1h |
| *SCR1* | scro1 | GRMZM2G131516 | Zm00001d052380_T001 | Zm00001eb195650 | Hughes et al. 2019* | AtSCR | |
| *SCR1h* | Gras 37 | GRMZM2G015080 | Zm00001d005029_T001 | Zm00001eb093670 | Hughes et al. 2019* | AtSCR | |
| *SHR1* | Gras 58 | GRMZM2G132794 | Zm00001d029607_T001 | Zm00001eb020840 | Schuler et al. 2017* | AtSHR; OsSHR2 | |
| *SHR2* | Gras 19 | GRMZM2G172657 | Zm00001d021973_T001 | Zm00001eb326020 | Schuler et al. 2017* | AtSHR; OsSHR1 | |
| *SHR2h* | Gras 85 | GRMZM2G019060 | Zm00001d006721_T001 | Zm00001eb108090 | Schuler et al. 2017* | AtSHR; OsSHR1 | |
| *RVN1* | IDD10 | GRMZM2G143723 | Zm00001d039467_T001 | Zm00001eb121010 | Sedelnikova et al. 2018 | – | |
| *RVN1h* | IDD 18 | GRMZM2G046290 | Zm00001d008842_T001 | Zm00001eb337700 | Sedelnikova et al. 2018 | – | |
| *ANT4* | EREB161 | GRMZM2G021573 | Zm00001d048004_T002 | Zm00001eb399440 | Liu et al. 2019* | ANT1-4 all co-orthologs of AtANT1 | |
| *PIN1a* | pin1 | GRMZM2G098643 | Zm00001d044812_T001 | Zm00001eb372180 | Matthes et al. 2019* | Co-ortholog AtPIN1, ortholog OsPIN1a | |
| *PIN1d* | pin4 | GRMZM2G171702 | Zm00001d052442_T001 | Zm00001eb196240 | O'Connor et al 2014* | Ortholog of OsPIN1c & OsPIN1d | Also named ZmPIN1d in many papers |
| *LAX1* | aic3 | GRMZM2G149481 | Zm00001d028401_T005 | Zm00001eb010460 | Matthes et al. 2019* | Co-ortholog AtLAX2, OsLAX2 & OsLAX4 | Named ZmLAX2 in Liu et al 2022 |
| *LAX2* | aic1 | GRMZM2G129413 | Zm00001d030310_T001 | Zm00001eb026490 | Matthes et al. 2019* | Co-ortholog AtLAX2, ortholog of OsLAX4 | |
| *LAX3* | atl1 | GRMZM2G127949 | Zm00001d042809_T003 | Zm00001eb147480 | Matthes et al. 2019* | Co-ortholog AtAUX1 & AtLAX1, ortholog of OsLAX1 | |
| *LAX5* | aic4 | GRMZM2G067022 | Zm00001d038275_T005 | Zm00001eb288800 | Matthes et al. 2019* | Co-ortholog AtAUX1 & AtLAX1, ortholog of OsLAX3 | |
| *HB52* | hb52 | –( AC187157.4_FG005) | Zm00001d008869_T001 | Zm00001eb337970 | TAIR | Ortholog of AtHB8 & AtHB15 | |
| *ARF4* | arftf4 | GRMZM2G034840 | Zm00001d001945_T010 | Zm00001eb067270 | Matthes et al. 2019* | Co-ortholog of AtMP, OsARF11 | |
| *ARF29* | arftf29 | GRMZM2G086949 | Zm00001d026540_T016 | Zm00001eb433020 | Matthes et al. 2019* | Co-ortholog of AtMP, OsARF11 | |
| *ARF16* | arftf16 | GRMZM2G028980 | Zm00001d053819_T008 | Zm00001eb207450 | Matthes et al. 2019* | OsARF6 | |
| *ARF34* | arftf34 | GRMZM2G081158 | Zm00001d031064_T006 | Zm00001eb031700 | Matthes et al. 2019* | Co-ortholog of AtARF6, ortholog of OsARF17 | |
| *ARF23* | arftf23 | GRMZM2G441325 | Zm00001d038698_T007 | Zm00001eb292830 | Matthes et al. 2019* | Co-ortholog of AtARF3 & AtARF4, ortholog of OsARF14 | |
| *KN1* | kn1 | GRMZM2G017087 | Zm00001d033859_T002 | Zm00001eb055920 | Hake et al. 1989 | Class I KNOX | |
| *MYB67* | myb67 | GRMZM2G089244 | Zm00001d053060_T001 | Zm00001eb201150 | Maize GDB | – | |
| *CYC4* | cyc4 | GRMZM2G310115 | Zm00001d012560_T002 | Zm00001eb369340 | Maize GDB | Ortholog of AtCYCB1 genes | cyc7 in raw data |
| *CYC7* | cyc7 | GRMZM2G073003 | Zm00001d043164_T001 | Zm00001eb150450 | Maize GDB | Ortholog of AtCYCB1 genes | previously cyc4b, cyc4 in raw data |
| *CYC3* | cyc3 | GRMZM2G073671 | Zm00001d036360_T002 | Zm00001eb272010 | Maize GDB | Ortholog AtCYCB2 genes | |

**Table S2. Gene expression in P0 cells identified from the 'peripheral zone' cluster.**

| Gene.ID | E1B2_M_P1_23 | E1B2_M_P1_121 | E1B2_M_P1_191 | S1A1_M_P1_39 | S1A1_M_P1_49 | S1A1_M_P1_82 | S1A1_M_P1_83 | **S1A1_M_P1_88** | **S1A1_M_P1_95** | **S1A1_M_P1_100** | **S1A1_M_P1_124** | **S1A1_M_P1_127** | **S1A1_M_P1_150** | S1A1_M_P1_160 | S1A2_M_P1_16 | S1A2_M_P1_72 | S1A2_M_P1_81 | S1A2_M_P1_234 | S1B1_M_P1_183 |
|---|---|---|---|---|---|---|---|---|---|---|---|---|---|---|---|---|---|---|---|
| SHR2 | 0 | 0 | 0 | 0 | 0 | 0 | 0 | 0 | 0 | 0 | 0 | 0 | 0 | 0 | 0 | 0 | 0 | 0 | 0 |
| TML2 | 0 | 0 | 0 | 0 | 0 | 0 | 0 | 0 | 0 | 0 | 0 | 0 | 0 | 0 | 0 | 0 | 0 | 0 | 0 |
| PIN1a | 0 | 0 | 1 | 0 | 0 | 3 | 0 | 1 | 3 | 6 | 0 | 0 | 3 | 0 | 1 | 0 | 0 | 0 | 0 |
| SCR1h | 0 | 0 | 0 | 0 | 1 | 1 | 0 | 0 | 0 | 0 | 0 | 0 | 0 | 0 | 0 | 0 | 0 | 0 | 1 |
| RVN1 | 0 | 0 | 0 | 0 | 0 | 0 | 0 | 0 | 0 | 0 | 0 | 0 | 0 | 0 | 0 | 0 | 0 | 0 | 0 |
| KN1 | 1 | 3 | 0 | 0 | 0 | 14 | 0 | 0 | 0 | 0 | 0 | 0 | 0 | 0 | 0 | 0 | 0 | 0 | 0 |
| CYC3 | 1 | 0 | 0 | 0 | 0 | 0 | 0 | 0 | 1 | 2 | 0 | 1 | 0 | 0 | 0 | 0 | 0 | 0 | 0 |
| CYC4 | 0 | 0 | 0 | 0 | 0 | 0 | 0 | 0 | 0 | 5 | 2 | 0 | 0 | 0 | 0 | 0 | 0 | 0 | 0 |
| ANT4 | 0 | 0 | 0 | 2 | 0 | 0 | 3 | 0 | 0 | 5 | 1 | 0 | 4 | 6 | 0 | 0 | 2 | 0 | 0 |
| PIN1d | 0 | 0 | 0 | 0 | 2 | 0 | 7 | 6 | 5 | 16 | 2 | 3 | 10 | 4 | 0 | 2 | 1 | 3 | 1 |
| LAX1 | 0 | 0 | 0 | 0 | 0 | 0 | 0 | 1 | 0 | 1 | 0 | 2 | 0 | 0 | 0 | 0 | 1 | 0 | 1 |
| LAX3 | 0 | 0 | 1 | 1 | 0 | 8 | 2 | 7 | 1 | 9 | 2 | 3 | 8 | 0 | 0 | 1 | 2 | 2 | 0 |
| LAX5 | 1 | 1 | 1 | 1 | 0 | 2 | 1 | 2 | 2 | 5 | 6 | 3 | 5 | 2 | 1 | 1 | 1 | 1 | 1 |
| HB52 | 0 | 1 | 0 | 0 | 2 | 1 | 0 | 3 | 1 | 5 | 4 | 4 | 2 | 7 | 11 | 4 | 5 | 6 | 0 |
| ARF4 | 1 | 1 | 3 | 0 | 0 | 7 | 4 | 1 | 2 | 0 | 5 | 4 | 0 | 0 | 3 | 1 | 1 | 6 | 0 |
| ARF29 | 1 | 0 | 0 | 0 | 0 | 0 | 0 | 0 | 0 | 0 | 0 | 0 | 0 | 0 | 0 | 0 | 0 | 0 | 0 |
| ARF16 | 0 | 0 | 0 | 0 | 3 | 8 | 2 | 5 | 4 | 2 | 5 | 6 | 1 | 6 | 0 | 0 | 0 | 5 | 1 |
| ARF34 | 5 | 4 | 3 | 2 | 7 | 0 | 1 | 7 | 5 | 9 | 4 | 11 | 7 | 5 | 2 | 6 | 3 | 1 | 5 |
| ARF23 | 0 | 0 | 0 | 0 | 6 | 1 | 3 | 3 | 10 | 21 | 12 | 14 | 13 | 3 | 1 | 0 | 4 | 2 | 1 |
| MYB67 | 0 | 0 | 0 | 0 | 0 | 0 | 0 | 1 | 0 | 0 | 0 | 1 | 1 | 0 | 0 | 0 | 0 | 0 | 0 |
| CYC7 | 1 | 0 | 1 | 1 | 7 | 0 | 0 | 3 | 15 | 21 | 27 | 14 | 2 | 3 | 9 | 0 | 1 | 0 | 0 |

**Table S3. Lateral and intermediate vein width quantification across the medio-lateral axis of leaves three and four.**

| LEAF 3 | | | | Lateral veins width ($\mu$m) - half leaf | | | | # Intermediates - half leaf | | | | Avg. intermediate veins width ($\mu$m) - half leaf | | | |
|---|---|---|---|---|---|---|---|---|---|---|---|---|---|---|---|
| Plant ID | Blade width (mm) | Blade length (mm) | # Laterals - half leaf | L1 | L2 | L3 | L4 | L1-L2 | L2-L3 | L3-L4 | L4-margin | L1-L2 | L2-L3 | L3-L4 | L4-margin |
| 1 | 14.49 | 330 | 4 | 112.89 | 106 | 88.24 | 89.64 | 15 | 14 | 8 | 5 | 48.9 | 45 | 45.3 | 44.4 |
| 2 | 16.56 | 360 | 4 | 118.44 | 121.82 | 97.7 | 83.81 | 16 | 16 | 8 | 4 | 49.75 | 49.36 | 42.67 | 41.21 |
| 3 | 16.8 | 365 | 4 | 150.73 | 116.4 | 106.87 | 76.53 | 12 | 14 | 13 | 11 | 52.42 | 48.28 | 46.69 | 49.73 |
| 4 | 17.99 | 300 | 4 | 138.18 | 131.45 | 125.36 | 83.93 | 13 | 14 | 12 | 6 | 62.4 | 53.77 | 43.29 | 43.53 |
| 5 | 14.79 | 310 | 4 | 110.92 | 105.63 | 95.02 | 83.19 | 14 | 15 | 10 | 4 | 53.06 | 42.79 | 35.22 | 44.29 |
| **Avg.** | **16.13** | **333.00** | **4.00** | **126.23** | **116.26** | **102.64** | **83.42** | **14.00** | **14.60** | **10.20** | **6.00** | **53.31** | **47.84** | **42.63** | **44.63** |
| stdev | 1.46 | 29.07 | 0.00 | 17.44 | 10.95 | 14.35 | 4.65 | 1.58 | 0.89 | 2.28 | 2.92 | 5.38 | 4.22 | 4.44 | 3.13 |

| LEAF 4 | | | | Lateral veins width ($\mu$m) - half leaf | | | | | # Intermediates - half leaf | | | | | Avg. intermediate veins width ($\mu$m) - half leaf | | | | |
|---|---|---|---|---|---|---|---|---|---|---|---|---|---|---|---|---|---|---|
| Plant ID | Blade width (mm) | Blade length (mm) | # Laterals - half leaf | L1 | L2 | L3 | L4 | L5 | L1-L2 | L2-L3 | L3-L4 | L4-L5 | L5-margin | L1-L2 | L2-L3 | L3-L4 | L4-L5 | L5-margin |
| 1 | 20.33 | 460 | 5 | 112.3 | 140.05 | 124.7 | 107.75 | 87.82 | 14 | 16 | 14 | 8 | 9 | 49.4 | 43.89 | 48.77 | 57.43 | 51.09 |
| 2 | 24.1 | 460 | 5 | 147.23 | 153.06 | 137.04 | 120.53 | 82.18 | 14 | 15 | 13 | 8 | 4 | 53.11 | 49.98 | 51.27 | 48.67 | 56.02 |
| 3 | 23.19 | 470 | 5 | 137.34 | 122.93 | 125.94 | 103.3 | 83.93 | 15 | 15 | 14 | 11 | 9 | 47.86 | 51.97 | 52.78 | 48.59 | 44.51 |
| 4 | 20.89 | 440 | 5 | 118.7 | 128.56 | 153.13 | 115.5 | 89.7 | 15 | 17 | 17 | 8 | 5 | 48.58 | 57.1 | 54.66 | 45.65 | 55.88 |
| 5 | 22.33 | 420 | 5 | 120.1 | 127.34 | 130.81 | 98.77 | 101.37 | 10 | 12 | 14 | 10 | 7 | 65.58 | 56.91 | 55.98 | 58.68 | 45.7 |
| **Avg.** | **22.17** | **450.00** | **5.00** | **127.13** | **134.39** | **134.32** | **109.17** | **89.00** | **13.60** | **15.00** | **14.40** | **9.00** | **6.80** | **52.91** | **51.97** | **52.69** | **51.80** | **50.64** |
| stdev | 1.57 | 20.00 | 0.00 | 14.57 | 12.20 | 11.58 | 8.86 | 7.54 | 2.07 | 1.87 | 1.52 | 1.41 | 2.28 | 7.37 | 5.48 | 2.83 | 5.85 | 5.44 |

**Dataset S1 (separate file).** Unprocessed and processed (normalised and scaled) transcript counts for stages M-P1, P2, P3, P4. Cells are labelled based on the section of origin (E1B2, E1D2, S1A1, S1A2, S1B1 or S1B2), and the plastochron stage under analysis (M-P1, P2, P3, P4). These data were used to generate Figs. 2, 4 and 5.

**Dataset S2 (separate file).** Molecular Cartography data clustering. Identities are labelled as outlined in Figs. 2A, 4A, 5A-B and S4. Cell labels as in Data S1.

**Dataset S3 (separate file).** Unprocessed transcript counts for the analysis shown in Fig. S5.

**Dataset S4 (separate file).** Normalised and scaled data obtained from Data S3. Refers to the analysis shown in Fig. S5.
